# Weighted Ensemble Simulations Reveal Novel Conformations and Modulator Effects in Hepatitis B Virus Capsid Assembly

**DOI:** 10.1101/2025.03.15.643452

**Authors:** Diane L. Lynch, Zixing Fan, Anna Pavlova, James C. Gumbart

## Abstract

Molecular dynamics (MD) simulations provide a detailed description of biophysical processes allowing mechanistic questions to be addressed at the atomic level. The promise of such approaches is partly hampered by well known sampling issues of typical simulations, where time scales available are significantly shorter than the process of interest. For the system of interest here, the binding of modulators of Hepatitis B virus capsid self-assembly, the binding site is at a flexible protein-protein interface. Characterization of the conformational landscape and how it is altered upon ligand binding is thus a prerequisite for a complete mechanistic description of capsid assembly modulation. However, such a description can be difficult due to the aforementioned sampling issues of standard MD, and enhanced sampling strategies are required. Here we employ the Weighted Ensemble methodology to characterize the free-energy landscape of our earlier determined functionally relevant progress coordinates. It is shown that this approach provides conformations outside those sampled by standard MD, as well as an increased number of structures with correspondingly enlarged binding pockets conducive to ligand binding, illustrating the utility of Weighted Ensemble for computational drug development.

**TOC Graphic:** 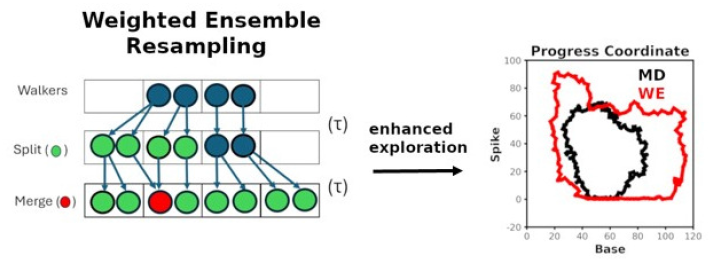

## Introduction

Significant progress has been made in experimental approaches to resolve the underlying dynamics for proteins, for example solid-state nuclear magnetic resonance (NMR),^1^ HDX-MS,^2^ and time-resolved cryo-EM ^3^ among others; however, transient states often remain experimentally inaccessible. While molecular dynamics (MD) simulations in principle can fully describe the dynamics, typical all-atom MD often can not reach physiologically relevant timescales. For example, it is only with specialized hardware that Ayaz et al.^4^ observe both ligand binding and resulting conformational changes MD simulations employing hundreds of *µ*s of simulation.

Hepatitis B virus (HBV) infection leads to severe liver damage and is the leading cause of liver disease^5^ with approximately a million deaths per year.^6^ HBV is an enveloped nucleocapsid, including an icosahedral shell structure comprising 120 core protein (Cp) homodimers (Fig. 1) containing the viral genome, which is surrounded by a membrane envelope.^7^ The capsid contains, protects, and delivers the viral genome during the HBV life cycle. Recent work has shown that a promising approach for reducing viral replication is to target the HBV nucleocapsid with ligands that disrupt capsid self-assembly (capsid assembly modulators or CAMs) producing non-infectious particles.^8,9^ The continued development of CAMs is an active area of research.^6^

**Figure 1:**
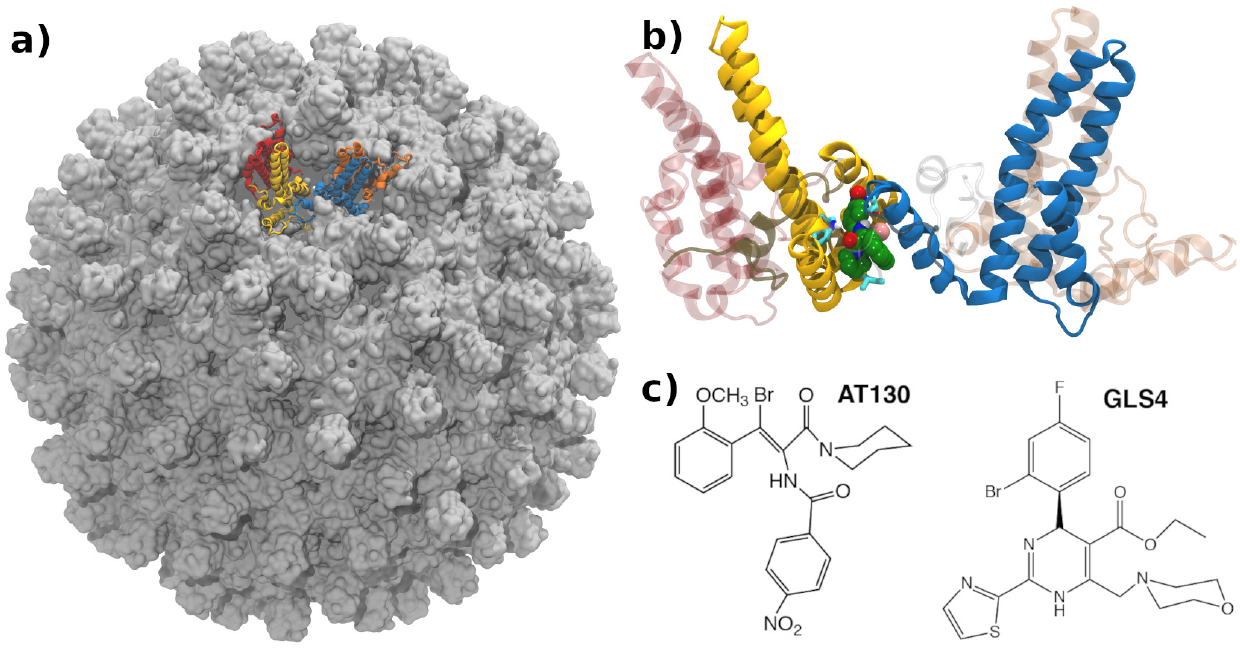
HBV capsid structure (PDB: 6HU4). a) The 240-unit capsid structure contains four quasi-equivalent tetramers. The ABCD tetramer, used for earlier MD studies, ^14^ is rendered in ribbon with the A, B, C, D subunits colored in red, gold, blue, and dark orange, respectively. b)The tetrameric unit with a bound CAM shown in a space-filling representation. c) Chemical structures of AT-130 and GLS4.

The building blocks of HBV capsids are core protein (Cp) dimers, which oligomerize into early-assembly intermediates, e.g., tetramers and hexamers, and ultimately the full capsid.^10,11^ Formation of the hexamer via the tetramer is the rate-limiting step of assembly.^10–12^ These earlier studies,^10,11^ supported by MD simulations,^13,14^ posit that the dimer’s conformation must undergo a shift to an “assembly active” state for productive assembly, as such capsid formation is described by a conformational selection mechanism. Environmental and other factors, such as CAMs, certain mutations, or ion concentration can alter the conformational landscape, resulting in capsid assembly modulation.^10–12,15–19^

Currently CAMs are distinguished based on their effects on capsid formation, where class II compounds are those that accelerate the production of normal, yet empty capsids^18,19^ (phenylpropenamides, e.g., AT-130), while class I CAMs generate aberrant structures such as tubes and sheets^17,20^ (e.g., heteroaryldihydropyrimidines like GLS4). Although assembly outcomes are significantly different, surprisingly CAMs share the same binding site at the tetrameric inter-dimer interface (Fig. 1b, c). Mechanistically this points to allosteric effects, ^21^ with CAM-specific alterations in the tetramer conformational ensemble, although rational control of such effects remain elusive.

Given the allosteric nature of HBV capsid assembly modulation, a detailed mechanistic description for capsid assembly will require a comprehensive evaluation of CAM specific modulations of the conformational ensemble. Moreover, comprehensive docking efforts in structure based drug design (SBDD) programs commonly employ ensemble docking, where protein flexibility is approximated by using a collection of diverse protein structures. ^22^ However, the success of such endeavors is dependent on the structures comprising these collections, which can be limited in the conformational space explored in standard MD simulations.

Various approaches to more fully explore the free energy landscape have been employed in MD applications, such as metadynamics, replica-exchange molecular dynamics, or other enhanced sampling approaches.^23^ In addition, a straightforward approach to accelerate sampling is to employ hydrogen mass repartitioning (HMR),^24^ allowing a larger (4-fs) time step for MD simulations. Earlier reports have established the validity of HMR in the equilibrium sampling of proteins^24^ as well as membrane systems,^25^ although the use of HMR for proteinligand kinetic quantities has recently been questioned.^26^ Intriguingly, the weighted ensemble (WE) simulation approach^27^ has been developed for rare-event processes and rigorously describes kinetic events, although it has also been employed to effectively generate an equilibrium ensemble.^28,29^ WE simulations offer the benefit of both enhanced sampling as well as unbiased dynamics and has been applied to a variety of systems, including *k*_on_ and *k*_off_ rates of protein-protein or protein-ligand binding,^30–32^ protein folding events,^33^ conformational changes,^34,35^ drug-membrane partitioning,^36^ as well as phase separation^37^ and continues to be developed with an open source python-based implementation readily available.^38^ Moreover, Xu et al.^39^ and Hellemann and Durrant^40^ have recently applied WE simulations for the study of ligand binding site properties in the context of ensemble docking studies.

Here, we have employed WE simulations, coupled with HMR, to explore the conformational space of apo, as well as CAM-bound, HBV tetramers. We demonstrate that WE-generated apo structures are capable of binding large HBV CAM ligands, with an increase in the number of conformations with large-volume ligand binding sites. Moreover, given the recent report of difficulties of HMR with ligand-based systems,^26^ we report a comparison of standard time-step MD with HMR for apo and CAM bound (AT-130 and GLS4; Fig. 1c) tetrameric systems. Overall, we find that the use of HMR in equilibrium simulations does not significantly perturb these protein-ligand systems and that WE simulations provide an effective approach for producing an expanded representation of the conformational ensemble, including a collection of unique structures that are difficult to access via standard MD.

## Results and Discussion

### HMR produces only modest effects on the conformational sampling of the HBV tetramer

In the course of our work, Sahil et al.^26^ reported sensitivity of ligand unbinding kinetics to the use of HMR. Their study illustrated that the use of HMR produces slowed ligand binding events, attributed to the effects of the longer timestep on both ligand diffusion, an effect observed earlier,^25^ and protein fluctuations. Here we have studied apo, AT-130, and GLS4 bound HBV tetramers using both a standard 2-fs time-step as well as HMR (4-fs time step) in order to characterize the distributions observed and point to any potential artifacts introduced via the application of the longer HMR time step. Using 12 replicas for each system, each simulation lasting 500 ns, root mean square deviation (RMSD) to the starting configuration, as well as the pairwise (all to all) RMSDs, which characterize the conformational space explored, were computed. In addition for both ligand-bound simulations after protein superimposition the RMSD, using the CAM heavy atoms, were calculated.f

On average, the 2-fs and HMR results display similar behavior for the systems studied, i.e., apo, AT-130 bound, and GLS4 bound. The average RMSD computed using the 2-fs and 4-fs trajectories compared to the starting structure for protein, CAM binding pocket, and ligand are within standard error (Figure S1). Although the RMSD distributions for individual replicas appear noisy (Figure S2), the overall protein distributions are quite similar (Figure S2a), while the analogous distributions for the CAM binding site using the 4-fs data appears shifted and somewhat wider than the 2-fs results (Figure S2d). Pairwise RMSDs (Figures S3 and S4) for the protein and CAM binding site were also computed, with the pair-wise computation measuring RMSD over the entire set of frames, rather than deviation from just the initial structure. Again, although the individual replicas appear noisy, the overall protein distributions are quite similar (Figure S4a), while the CAM binding pocket in the 4-fs trajectories appears variable and slightly shifted (Figure S4d-f). Analogous analysis was performed for the ligand bound AT-130 and GLS4 simulations (Figures S5-S10) with similar results to those observed for the apo simulations, i.e., the introduction of HMR produces very small alterations in the protein structures, while the CAM binding site appears more susceptible to noise. However, it is important to note that convergence of such simulations can be difficult, with multiple replicas recommended with the number dependent on the length of the trajectory as well as the system under study.^41^ Given the goal in the present study is the exploration of the equilibrium conformational ensemble, rather than kinetic properties^26^ of the HBV tetramer, we have used HMR in the following WE simulations.

### Effects of CAM binding on tetrameric structures

Our biophysics-based approach^14,42^ for exploring the molecular details of CAM specific capsid assembly modulation has revealed CAM-class specific altered tetrameric conformations, which are associated with accelerated, yet normal (Class II), or aberrant capsid (Class I) assembly. Upon CAM binding, ligand-specific structural changes of the HBV dimer occur, which generate intermediates that ultimately lead to altered capsid assembly. As such CAM binding is characterized by both its binding affinity as well as the degree to which it alters the underlying tetrameric conformational ensemble.

Pavlova et al.^14^ established that angles defining the inter-dimer orientation, i.e. the base and spike angles (Fig. 2a), are effective at discriminating the conformational preferences of the apo, as well as CAM-bound tetramers. The base and spike angles describe the opening and closing or bending of the tetramer respectively, with the base angle defined by the *α*5 helices of the Cp monomers and the spike angle defined by the relative direction of the *α*3/*α*4 helices in each dimer. From these earlier simulations it was apparent that the combination of these two angles characterized the observed ligand-dependent conformational variation in apo and CAM-bound simulations.^14,42^ Importantly these angles provide a mechanistically relevant reduced representation for describing the CAM specific modulation of the dynamics of the tetrameric inter-dimer orientations. Additionally, the use of HMR does not alter the average value of the base and spike angles (Figure S11).

**Figure 2:**
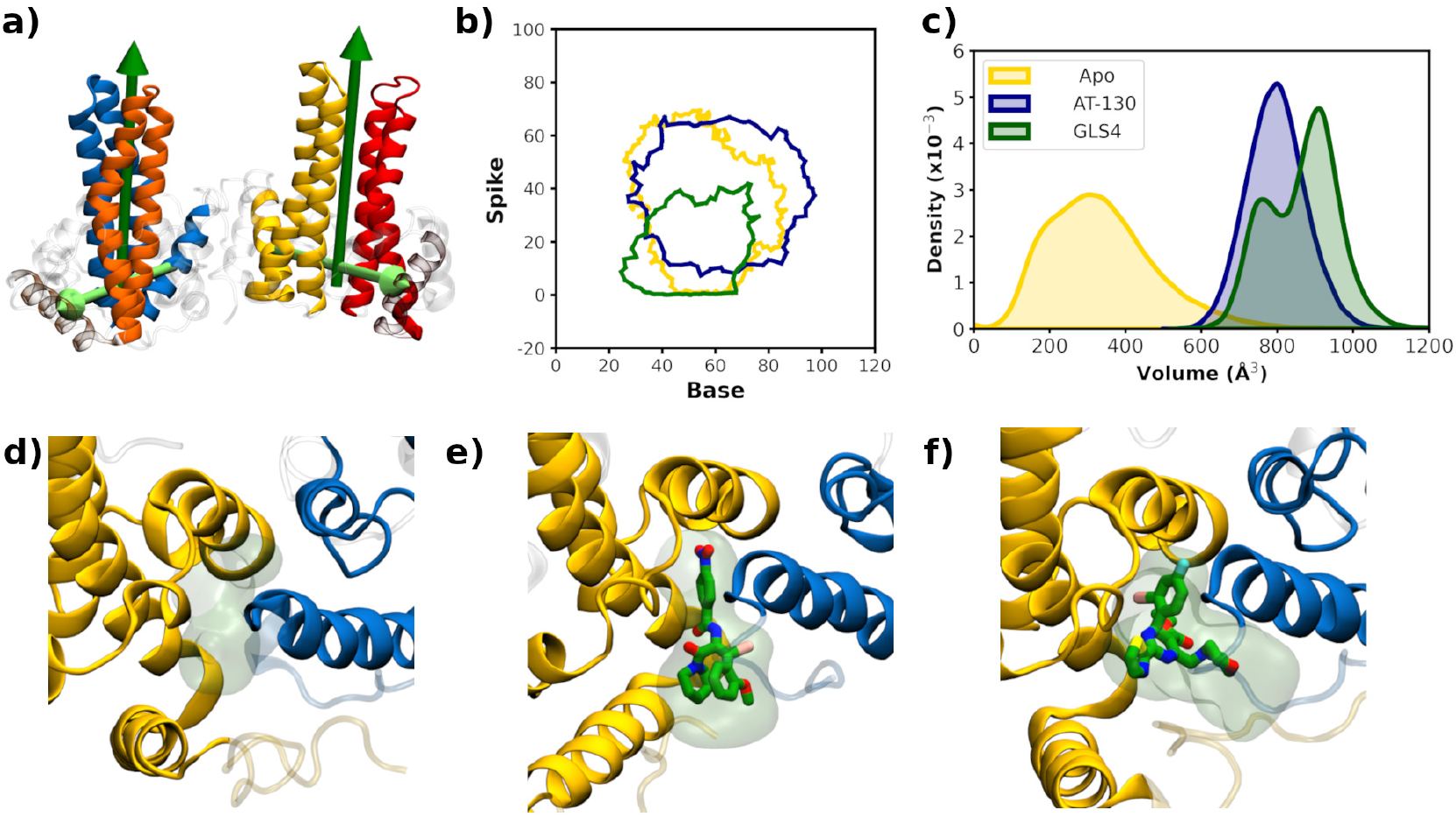
a) Illustration of the base and spike angles, with the vectors describing the base/spike angles rendered a light green/dark green, respectively. b) The base and spike angles for the apo (gold), AT-130 (dark blue), and GLS4 (dark green) rendered as a boundary for our standard MD simulations. c) Distribution of binding pocket volumes computed with Epock for the apo, AT-130, and GLS4 bound standard MD simulations. d)-f) Structures of the binding pocket volumes for the apo (volume = 287Å^2^), d) AT-130 (volume = 785Å^2^), and e) GLS4 (volume = 955Å^2^). Chain B is rendered in gold ribbon, chain C in dark blue ribbon, ligand is rendered in licorice with the carbons, oxygen, sulfur, or bromine atoms green, red, yellow, or pink. The outline of the Epock-derived volume is a transparent green surface.

### Standard MD simulations of the apo HBV tetramer produce limited exploration of base and spike angles

The HMR-based standard MD replicas described above, whose RMSD have all nearly reached a plateau (Figure S12), were used to evaluate the apo and CAM-specific base and spike angle landscape (Fig. 2b), with the angle data represented by a boundary encompassing all the data points. The class II misdirector (GLS4) base and spike angle pair are shifted to lower values and explore a more restricted region of inter-dimer orientations relative to those of the apo and AT-130 simulations, particularly the spike angle (average values and standard errors are reported in Figure S11). The apo and AT-130 results are clearly similar, and although noisy, the AT-130 simulations appear wider and reach larger base/spike values, reflecting its mechanism of action, namely assembly acceleration.

The binding site for CAM compounds, regardless of functional outcome, lies at the interface of the B and C chains in the tetrameric unit of the capsid (Fig. 1b). We have evaluated the CAM-binding-site volumes (see Methods for residues used) from the apo, AT-130, and GLS4 simulations (illustrated in Figures 2d-f) using Epock.^43^ The distributions observed in the apo simulations are spread over a large range and reflect the flexibility of the apo binding site (Fig. 2c), while the pocket is well defined in the AT-130- and GLS4-bound simulations. Relative to AT-130, the GLS4 ligand displaying a somewhat larger available volume. The secondary maximum at lower volume in the GLS4 simulations corresponds to a set of structures where the C-terminus extends around the ligand limiting available space (see Figure S13 for an illustration). Of particular note, very few structures from the apo simulation could accommodate either ligand, highlighting a known issue with using structures from apo simulations for docking.^44^

### WE simulations more effectively explore base and spike angles

Weighted ensemble simulations are a resampling strategy designed to produce rare-event transitions, without the addition of applied forces, along an appropriately chosen progress coordinate or coordinates^27^ and belong to the path-sampling class of enhanced sampling approaches. In addition to steady-state conditions, where an initial state is transitioned to a predefined target state, equilibrium conditions are also possible. ^28,45^ An advantage of the latter is that a target state definition is not necessary; as such, these simulations explore the conformational space of the system in an unbiased fashion.

Two replicas of WE simulations for the apo HBV tetramers were performed (6 µs total) employing 2D progress coordinates consisting of the base and spike angles discussed above. As detailed in the methods, each angle was partitioned into 20^?^ bins. The evolution of the base and spike angles are provided in Figure S14, while the 2D landscape generated from the combination of the two replicas is illustrated in Fig. 3a. It is apparent that although a single structure was employed to start the WE simulations, the equilibrium distribution spreads out such that a wide range of angles are sampled. In Fig. 3b and c, the weights as a function of iteration for two highly populated bins for the individual replicas are reported. Similar to the distribution of rate constants reported by Adhikari et al.,^33^ each replica behaves somewhat differently, although reasonable convergence of the highly populated bins is achieved. However, even for runs of 3 µs convergence of the smaller weights appears more problematic, e.g., the magnitude of the smaller weights in bins oscillates dramatically with small changes in population (Figure S15).

**Figure 3:**
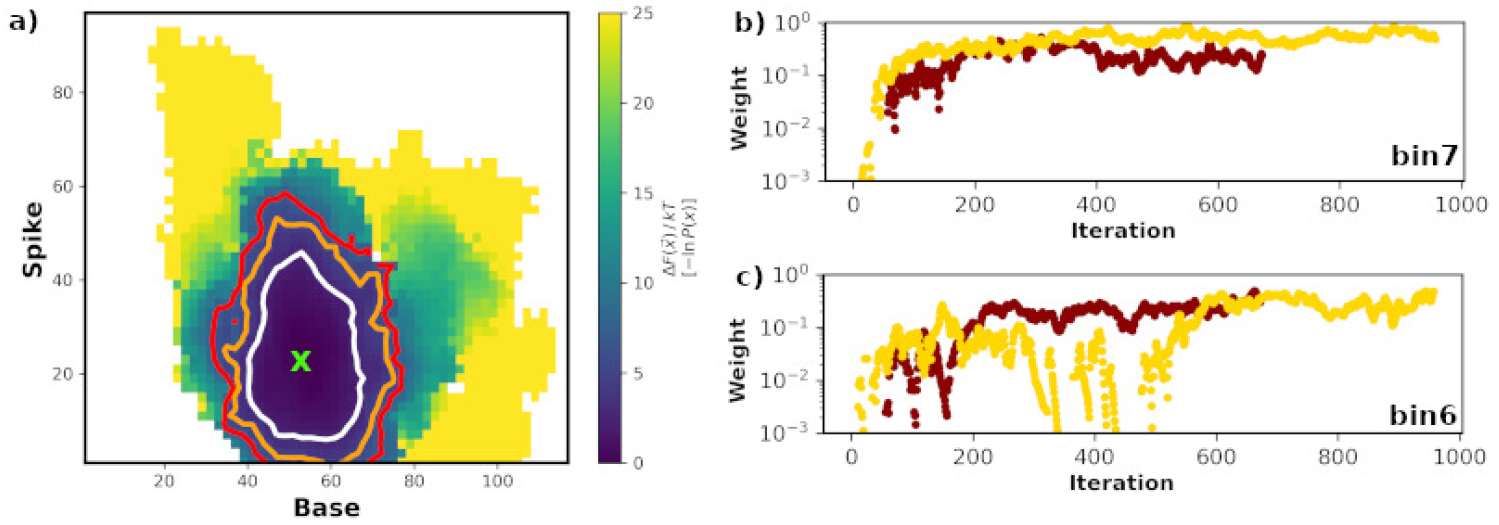
Apo WE simulations. a) 2D free energy landscape after combining the two replicas. Contoured at 2.5*kT* (white), 5.0*kT* (orange), and 7.5*kT* (red) and removing the first 250 iterations. The initial base/spike angle pair is indicated with a green X. b) and c) Bin weights as function of WE iteration for bins 6 (base=35°-55°, spike=30°-50°) and bin 7 (base=35°-55°, spike=30°-50°).

Analogous WE simulations, two replicas of 3 µs each, were performed for AT-130- and GLS4-bound tetramers, and, similar to the apo results, the major bins appear reasonably well converged. Figures S16-S19 display progress coordinate evolution, the 2D landscapes, and the weights as a function of iteration for these simulations. The resulting combined 2D landscapes for the CAM-bound simulations (Fig. 4) and progress coordinate evolution (Figures S16 and S18) reveal an altered exploration pattern for the GLS4-bound simulations relative to those for the apo and AT-130 bound systems. Similar to the apo WE results, the AT-130 bound tetramer explores a wide range of base and spike angles and is consistent with the shifting of these angles to larger values relative to the apo simulations (Fig. 3a) as observed in standard MD (Fig. 2b). Despite the fact that the current WE settings provide robust sampling along the base/spike progress coordinates for both apo and AT-130 simulations, in the case of GLS4, both replicas explore a much narrower range (Figure S18). Although these results are consistent with the observations that GLS4, a misdirector, does not readily sample the inter-dimer orientations necessary for normal capsid assembly,^14,42^ it is likely a more aggressive re-binning during the WE simulations, or the use of the adaptive binning procedure^46^ could provide further exploration of these inter-dimer orientations.

**Figure 4:**
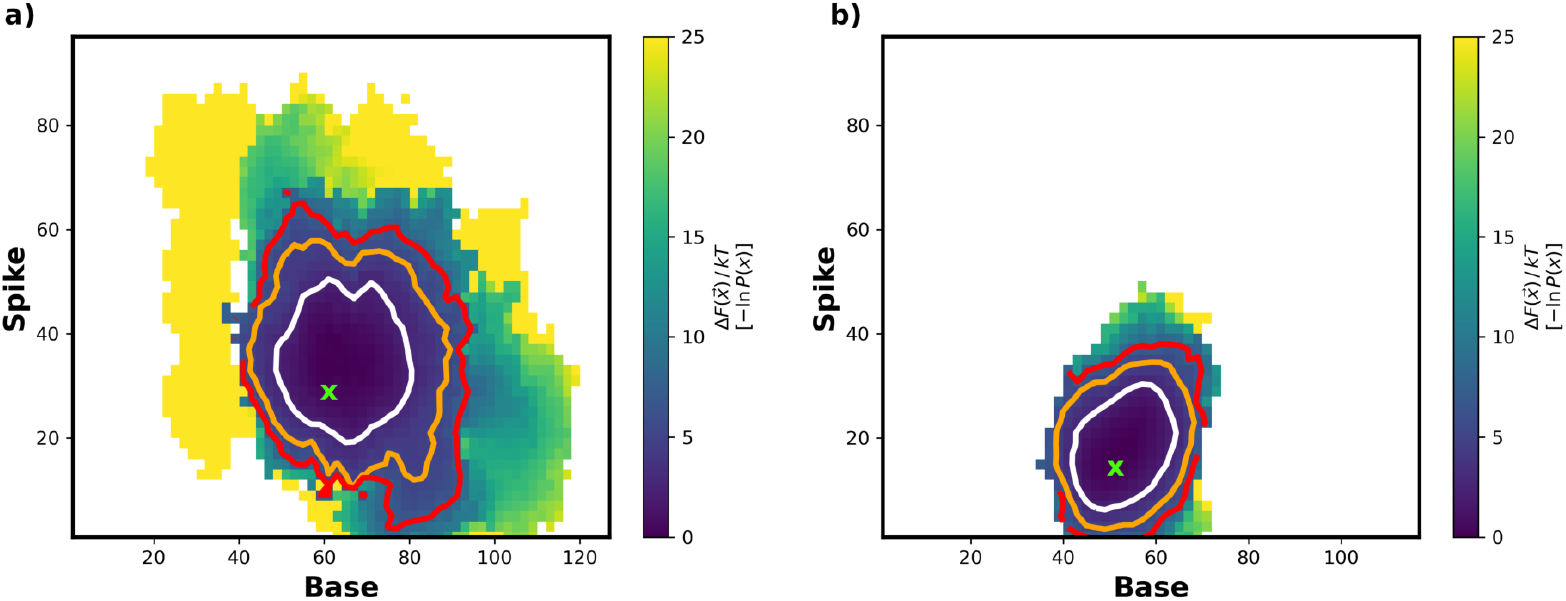
AT-130 and GLS4 WE 2D energy landscapes. a) AT-130 and b) GLS4 results after combining the two replicas (3 µs for each replica). Contoured at 2.5*kT* (white), 5.0*kT* (orange), and 7.5*kT* (red) and removing the first 250 iterations. The initial base and spike angle pair is indicated with a green X.

The WE conformational ensemble successfully produces additional conformations not observed in standard MD. To illustrate this point the base/spike region observed in the apo WE simulations is compared to standard MD in Figure 5a (both a collective 6 µs) where, as in (Fig. 2b) the standard MD results are represented via a boundary. Again, similar results are obtained for the AT-130 WE simulations, i.e., the WE results explore regions outside those sampled in standard MD (Figure S20a), while in contrast, the GLS4-bound tetramers in the WE simulations, relative to the standard MD results, sample a comparable region of the base/spike angle space (Figure S20c).

**Figure 5:**
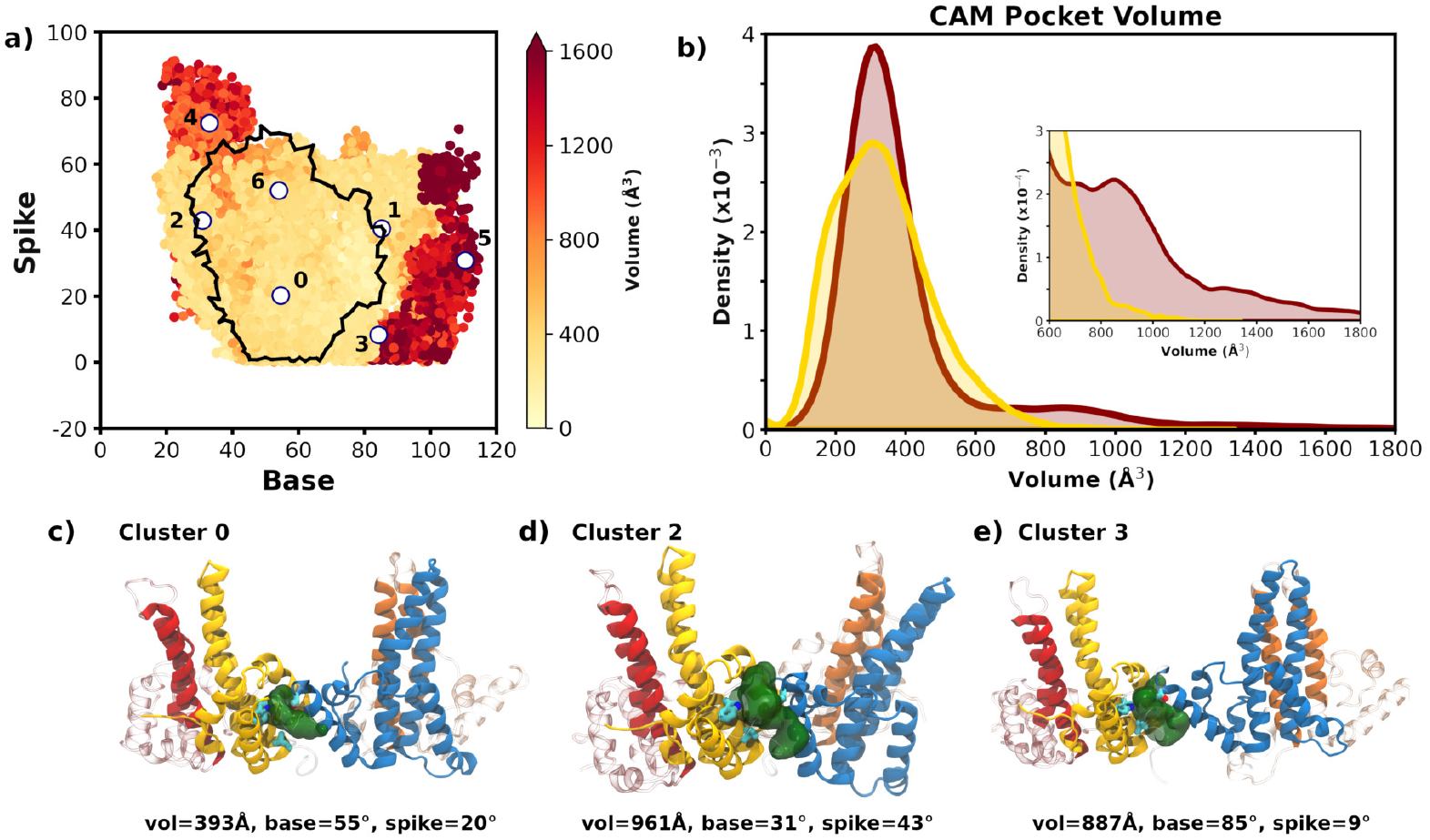
WE samples a wider distribution of base and spike angles. a) Scatter plot of the base and spike angle progress coordinates (unweighted). Collective data for the combined replicas is reported, with the first 250 iterations of WE simulation and the first 10 ns of standard MD data removed. For clarity, the standard MD data is rendered as a boundary (black line).The WE data is colored by the CAM pocket volume. Cluster centers are marked with white circles and are numbered. b) Histogram of the CAM pocket volumes for WE and standard MD, with the inset for the larger volumes. c) - e) Representative conformations of the tetramer structures from selected clusters showing the Epock pocket volume. W102, T128, and L140 are rendered in cyan licorice, and the volume rendered as a dark green surface.

### Apo WE simulations explore enlarged ligand binding sites

Holo structures are often preferred to their apo counterparts for drug discovery purposes as their ligand binding sites are already well-formed,^47^ although such structures may be unavailable for the protein of interest. In such cases, methods to address this shortcoming include template based approaches to binding site refinement,^48^ metadynamics to explore an enhanced sampling of the binding site,^44^ or induced-fit procedures.^49^ However, WE-based simulations are an attractive alternative to these approaches since the underlying dynamics remains unbiased.

Although we are focused on the CAM-specific allosteric modulation of the HBV tetramer, initial binding is still required. Therefore Epock^43^ was used to estimate the CAM pocket volumes for the WE-generated apo conformations. In Figure 5a, the unweighted WE data is colored by pocket volume; it is noteworthy that the larger volumes correspond to the more extreme values of the progress coordinates, whereas standard MD produces few, if any, conformations with larger volumes. Figure 5b displays the CAM pocket volume distribution for the WE replicas compared with pocket volumes from the standard MD simulations. The distributions are centered about a volume of ∼350 Å^3^ for both the WE and standard MD runs. Although the fraction of the total number of conformations with large pocket volumes is small relative to the standard MD results, WE simulations generate an increased number of conformations with larger-than-average pocket volume (Figure 5b, inset). These structures are likely to be amenable to docking larger CAMs than those with an average ligand binding site volume. We emphasize that the progress coordinates currently used do not explicitly include information concerning the ligand binding site; larger binding site volumes are naturally produced in the course of sampling the functionally relevant base and spike angles. Given that the number of conformations needed to generate a successful ensemble docking protocol can be quite large,^50^ our results suggest that the use of WE simulations will enhance the success rate in ensemble docking efforts. In addition to exploring the CAM binding pocket volume in the apo simulations, the AT-130 and GLS4 simulations were also analyzed (Figure S20b and c). As expected the presence of the ligand reduces the variation in the binding site volumes for both AT-130 as well as GLS4.

Clustering is often employed in ensemble docking in order to extract a small representative subset of protein conformations and is performed on global protein RMSD^22^ or, alternatively, RMSD of the ligand binding site residues directly.^51^ In the present case, although the binding pocket volume is required to be of sufficient size to bind a ligand, it is the functional, i.e., allosteric response of the tetramer, that is thought to drive modulation of HBV capsid assembly.

Both the WE (unweighted data) and standard MD data were clustered using hierarchal clustering, employing the base and spike angles. An average euclidean linkage^52^ was used. Both the WE and standard MD results were downsampled, with ∼25000 structures retained at a 200-ps interval for each. As discussed by Hellemann and Durrant,^40^ such downsampling may skew the distribution due to the merging and splitting steps in the WE protocol. However, we find that such effects are modest in the base and spike angle distributions used here (Figure S22). The WE data generated seven clusters, whose centers are labeled in Figure 5a, and span the range of base/spike angles. Representative conformations extracted from each cluster produce structures with a range of binding pocket volumes. Several of these conformations are illustrated in Figure 5c-e, labeled with their volumes, base, and spike angles, while structures from all seven clusters, along with the hierarchal dendrogram, are displayed in (Figure S23). It is noteworthy that some of the conformations with very extreme values for the base and spike angles have extremely large volumes (e.g. cluster center 5) and appear nearly dissociated. Clustering of the apo standard MD produced four clusters (Figure S24).

### Apo WE simulations provide suitable conformations for CAM docking

In order to validate that the conformations obtained from the apo simulations are conducive to docking, we extracted structures from clusters 0, 1, 2, 3, and 6 of the WE simulations and conformations from the four standard MD clusters and docked GLS4 to each. The ligand binding site volumes for the extracted conformations are reported in Table 1. GLS4 was chosen for these docking studies since it requires a significantly larger volume than that observed in our standard MD simulations (Figure 2b), thus posing a significant challenge for these apo-derived structures. Clusters 4 and 5 from the WE simulations were eliminated from this initial docking calculation as they have extreme values for the base and spike angle pairs.

**Table 1:** Ligand Binding Pocket Volumes (Å^3^) for representative conformations.

| Cluster Number | standard MD | WE | standard MD (extreme) |
| --- | --- | --- | --- |
| 0 | 244 | 393 | 860 |
| 1 | 452 | 471 | 897 |
| 2 | 160 | 961 | 1099 |
| 3 | 295 | 887 | 702 |
| 6 | - | 469 | - |

The GLS4 ligand molecule was docked into a volume centered around the CAM binding pocket of the protein using AutoDock Vina. Not unexpectedly, due to their small binding pocket volume, all four structures taken from the standard MD clusters failed to support ligand binding, with the ligand docked outside the pocket in all poses. However, in the case of the structures taken from the WE simulations, several produced ligand bound states. An example from cluster 2 illustrates that the pose reproduces essential interactions, i.e., the Trp102 sidechain interacts with the pyrimidine group, the ethyl-ester moiety is sandwiched between Thr109 on chain B and Val124 on chain C, and the morpholino group is proximal to Thr128.

Although we observed an enhancement of large volume structures in the WE simulations (Figure 5b inset), standard MD does produce some structures that generate expanded ligand binding sites. Structures were extracted from each of the four standard MD apo clusters that have base and spike angles significantly different from their cluster average as well as enlarged pocket volumes. Figure S25 presents snapshots of these conformations and details of the selection criterion used. Volumes for these structures are also included in Table 1. Docking of GLS4 to one of the clusters produced a pose that supports GLS4 binding (Figure 6c), similar to both the initial GLS4-bound conformation and the WE-derived docked one (Figure 6a, b). After superposition of the binding site residues, GLS4 heavy atom RMSDs to the initial conformation for the WE/standard-MD-derived structures are 2.9 Å and 2.2 Å, respectively. Given that the docking scores for these poses are both favorable (*<* -8 kcal/mol), these results highlight that although rare, conformations produced by WE simulations and even standard MD can accommodate large ligands such as GLS4. However, it has been noted that the number of structures necessary to generate a successful docking campaign can be considerable, while only one suitable structure was obtained with standard MD. Therefore, we contend that WE offers a reliable way to maximize the number of conformations that can be successfully employed.^50^

**Figure 6:**
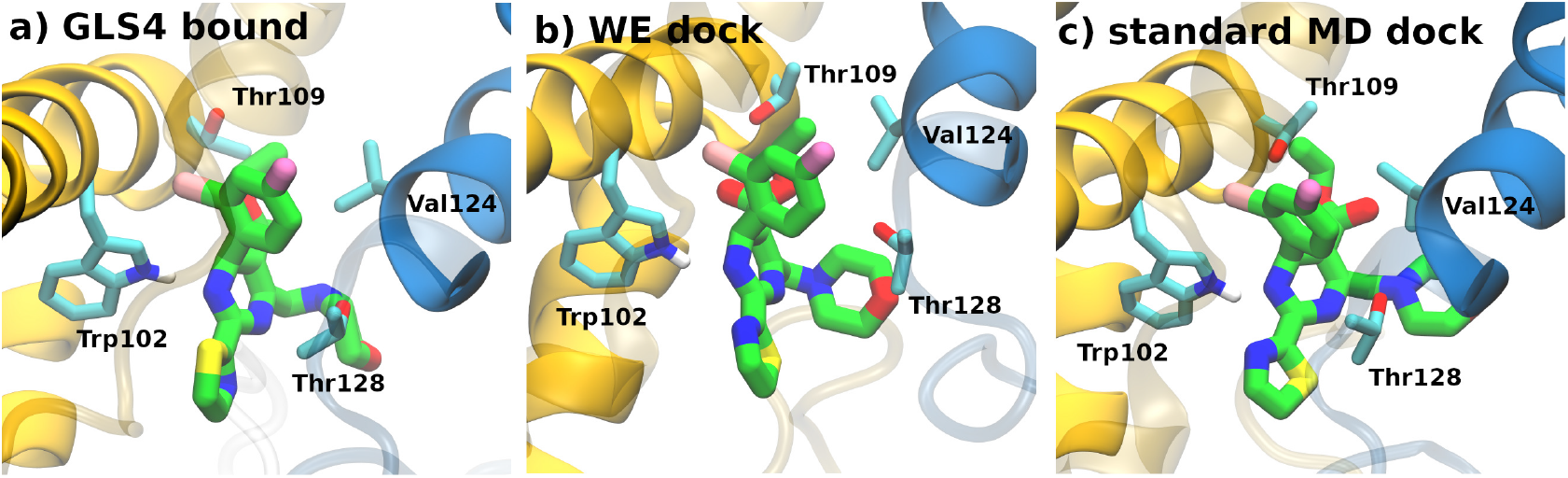
Docking of GLS4 to the tetramer. a) Initial conformation of the GLS4-bound structure. b)sample conformations taken from docking GLS4 into WE cluster 2 (docking score -9.3 kcal/mol) and c) structure taken from docking GLS4 into an extreme example from standard MD cluster 2 (docking score -8.3 kcal/mol).

## Conclusions

CAM efficacy is not only characterized by binding affinity but also by the resulting conformational changes in early-assembly intermediates such as the tetramer. Our study demonstrates that standard MD simulations for the apo HBV Cp tetramer rarely sample conformations that display ligand binding site volumes of sufficient size for CAM binding. Furthermore these standard MD simulations produce conformations which span a restricted region of the functionally relevant HBV tetramer inter-dimer orientations as characterized by the base and spike angles. However, both of these deficiencies are alleviated by the use of the Weighted Ensemble approach. The use of WE-based simulations not only produce conformations characterized by base and spike angles outside those sampled in standard MD, but also generate an enhanced number of larger volume CAM binding pockets. Furthermore, we show that these WE produced apo structures can be successfully used in docking GLS4, a reasonably large, well-studied CAM. These results open the way for further virtual screening studies targeting the allosteric modulation of HBV capsid assembly. Additionally, similar to results observed for the apo system, standard and WE-based MD simulations for the AT-130 bound case produce enhanced sampling of the HBV tetramer. In contrast, the exploration of the base and spike angle regions is comparable for the GLS4 bound state, suggesting a ligand-dependent effect. However, additional simulations are needed to confirm this. Finally, the use of HMR is shown to have only a modest effect on the protein and ligand equilibrium distributions. The present study, along with recent work by Xu et al.^39^ and Hellemann and Durrant,^40^ has established that the WE approach is an effective enhanced sampling method for generating and characterizing ligand binding alterations in the protein conformational landscape and produces viable conformations for further docking studies.

## Methods

### Molecular Dynamics

The starting structures for the apo, AT-130, and GLS4 simulations, based on the 3J2V,^53^ 4G93,^54^ and 5E0I^55^ (where the bound ligand was changed to GLS4) structures respectively, were taken from the previous equilibrated simulations of Pavlova et al.^14^ and minimized for 2000 steps prior to simulation. For standard molecular dynamics each system was run for 500 ns employing 12 replicas for an aggregate of 6 *µ*s of simulation. All MD simulations were carried out with NAMD3^56^ using the CHARMM36m force field for proteins^57^ along with the TIP3P water model.^58^ CHARMM force field parameters for GLS4 were taken from Pavlova et al.,^14^ where the CGenFF^59^ parameters via the webserver was used, while parameters for AT-130 were taken from Pang et al.^60^ For the standard MD simulations, either a 2-fs or a hydrogen mass repartitioning (HMR) scheme was employed, with the latter allowing the use of a 4-fs time step.^24,25^ In all simulations, the van der Waals cutoff was set to 1.2 nm, with a smoothing function applied from 1.0 to 1.2 nm. Long-range electrostatics were calculated using the particle mesh Ewald method.^61^ A constant temperature of 310 K was maintained using Langevin dynamics and a constant pressure of 1 atm was maintained using the Langevin piston method available in NAMD.^56^

Molecular dynamics settings for the evolution of the replicas in the WE simulations were identical to those of the standard HMR MD described above, while the initial configuration was taken after 40 ns of standard MD. WE specific parameters are described further below. Unless otherwise noted, visualization and analysis were done using VMD version 1.9.4.^62^ Pair-wise RMSD calculations were performed with MDTraj^63^ and delaunay triangluation, available in SciPy,^64^ was used for generating progress coordinate boundaries.

### Weighted Ensemble

In the current WE simulations a 2-D progress coordinate was chosen based on the results of earlier work^14^ which identified specific dimer-dimer orientations of the HBV tetramer, described by the base and spike angles of (Figure 2a), sensitive to the presence and/or type of CAM. The base angle reflects an opening or closing of the Cp dimers in the tetramer and is calculated based on the geometry of the constituent *α*-5 helices, defined using the backbone atoms of residues 111 to 127, specifically for each dimer the vector between the geometric centers of the two *α*-5 helices computed, and the angle between these 2 intra-dimer vectors is the base angle. The geometry of the *α*-3 and *α*-4 helices are used in the spike angle definition. Dividing these helices into an upper and lower portion about a flexible hinge generates two segments: an upper segment using residues 49 to 56 and residues 103 to 110 and a lower segment using residues 56 to 65 and 96 to 103. Given this definition the vector from the lower to upper center is evaluated for each dimer and the angle between these resultant vectors is the spike angle.

Although WE simulations are often run in order to provided unbiased kinetic date using steady state conditions, equilibrium conditions can also be applied. In fact WE has been used in order to effectively sample such equilibrium populations.^28,30^ Analogous to the initial/preparatory equilibrium simulations for peptide-protein and protein-protein interactions^29,33^ we employed equilibrium WE simulation using WESTPA version 1^65^ (release. 2020.06) using 2-D progress coordinates specified by the base and spike angles discussed above, until reasonable convergence was obtained for the larger population regions. The simulations were performed using equilibrium conditions, with a *τ* = 50ps, 8 trajectories per bin, and the bins subdivided at 20° boundaries for both the base and spike angles, initially set at [15,35,55,75,95] and [10,30,50,70,90] respectively. Given the variance observed in multiple WE runs^33^ and as recommended^66^ two independent WE simulations were performed, each run for a collective ∼3*µ*s each. In the case of the apo simulations this resulted in 650/960 iterations for each replica respectively, while for AT-130 and GLS4 bound simulations the iterations per replica are 610/500 and 1350/1365 respectively. Note, when the progress coordinate exceeded the extreme bin boundaries these were subsequently extended. The evolution of each progress coordinate and the 2-D landscape for each replica were generated using the WESTPA tools *w_pdist* and *plothist*, with the replicas combined using *w_multi west*.

### Epock Analysis

CAM pocket volumes were computed using Epock.^43^ The pocket was identified as those residues within a close contact (4.5 Å) between heavy atoms of the ligand in the AT-130 and GLS4 structures from the starting structures and include residues 121, 124, 125, 127, 128, 129, 132, 133, 134, 136, and 138 from chain C and residues 23, 25, 29, 30, 33, 37, 102, 105, 106, 109, 110, 114, 115, 118, 138, 139, and 140 from chain B. The volume was obtained using a spherical probe (10Å radii) along with non-contiguous points removal, which removes free space points outside a given radius here taken as 3 Å. Both are centered on the geometrical center of the binding site residues listed above. Additionally an exclusion region, defined by a 3 Å sphere centered at the COM of the sidechains of chain B residue 29 and chain C and residue 127, 129, and 133 was used. Binding pocket volumes using these choices are illustrated in Figure S26, where ligands under study are effectively encompassed.

### Clustering Analysis

The structures used for clustering were taken from the collection of all WE or standard MD conformations, the number of conformations used was 25412/27000 for the WE/standard MD respectively, nearly the same in order to cluster approximately the same amount of data at ∼0.2ns intervals. These differences result from the WE algorithm, which generates data after each iteration. In both cases initial data was dropped prior to analysis, 10ns in the case of standard MD and where the highly populated bin weights were approximately leveled off for the WE simulations, at 250 iterations. Xu et al.^39^ employed a different approach; with the WE simulations using ligand binding site information (SASA) to sample alternate conformations, with the clustering performed using a representative pathway taken from the WE simulation. In our case we have focused on the functionally important motion for HBV tetrameric assembly modulcation, i.e. extreme base and spike angle coordinates. In this case a representative trajectory may in fact miss the relevant apo conformations. Moreover, clustering WE simulations has been discussed by Hellemann and Durrant, where they employ a 2-tier approach to avoid striding over the branching points in a WE simulation.

The clustering of the base and spike angles was performed with SciPy^64^ hierarchical clustering, using average euclidean linkage, with the data preprocessed prior to clustering such that the data has zero mean and unit standard deviation using sklearn.^67^ The merging distance at each clustering step was used to determine the number of clusters, with a jump in the distance as another cluster pair is merged, indicating a separation in the data. Unless otherwise noted, i.e. for the apo standard MD described in Figure S25, the structure whose base and spike angles were closest to the center for each cluster was extracted for illustration and the docking calculations.

### Docking

The GLS4 molecule was docked into the capsid assembly modulator (CAM) binding pocket of the target protein using AutoDock Vina (version 1.1.2).^68,69^ To account for potential conformational adjustments during docking, the side chains of residues within a 6 Å radius of the GLS4 molecule were set as flexible. This flexibility ensures accurate modeling of protein-ligand interactions in the binding pocket.

The docking simulation explored a search space defined as a 30 Å × 30 Å × 30 Å box, centered on the original position of GLS4 within its bound conformation ((PDB code 5E0I). The center of this box was determined after structural alignment of the protein conformations to the GLS4-bound structure. To achieve robust and consistent docking results, the exhaustiveness parameter was set to its maximum allowable value of 32.

## Acknowledgement

This work was supported by the National Institutes of Health (R01-AI148740). We also used the Hive cluster in this work, which is supported by the National Science Foundation (MRI-1828187) and is managed by the Partnership for an Advanced Computing Environment at Georgia Tech.

