## Supporting Information for "Weighted Ensemble Simulations Reveal Novel Conformations and Modulator Effects in Hepatitis B Virus Capsid Assembly"

### Comparison of 2-fs and 4-fs(HMR) MD Simulations

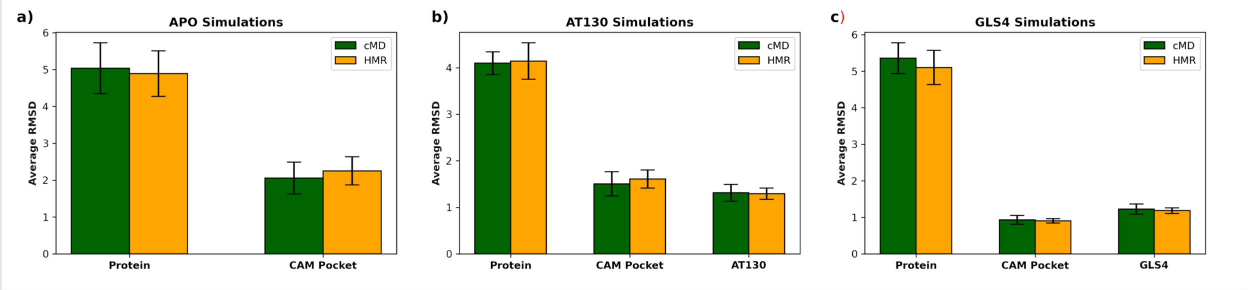


Figure S1: Average RMSD to the starting structure, with standard errors calculated from the 12 independant replicas, for the protein(aligned using all C$\alpha$ atoms), CAM binding pocket (aligned using the binding pocket C$\alpha$ atoms, defined in the Epock methods section), and ligands (heavy atoms). The RMSD of the ligand was computed after alignment of the CAM binding pocket residues.

### Apo HMR/2-fs Simulations

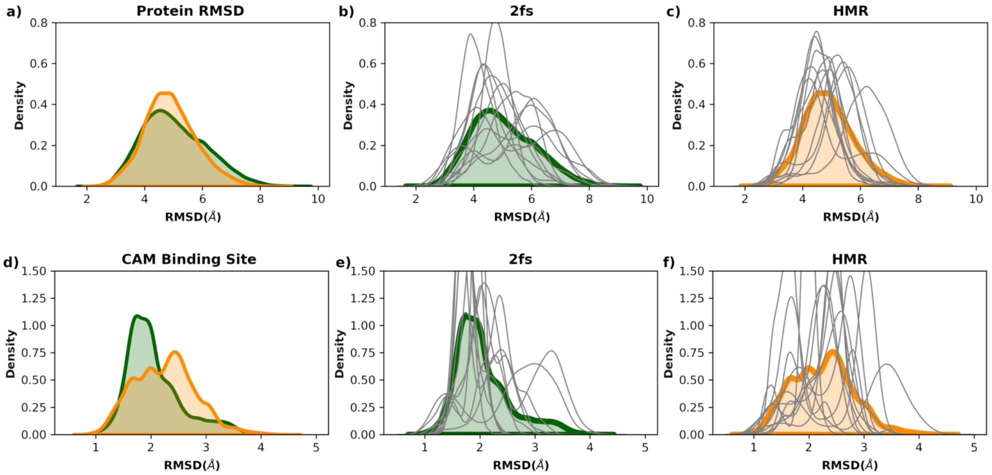


Figure S2: RMSD distributions to the starting structure from the apo simulations. Top row (panels a-c): RMSD aligned and calculated using all protein C$\alpha$ atoms. Bottom row (panels d-f): RMSD aligned and calculated using the C$\alpha$ atoms of the CAM binding site residues. Panels a) and d) report the combined replica RMSD distributions while the middle and right columns are the 2fs (panels b and e, with combined replicas: green lines) and HMR (panels c and f, with combined replicas: orange lines) respectively. The individual replica data is plotted separately (grey lines). Note, for clarity the replicas are separately normalized to 1.

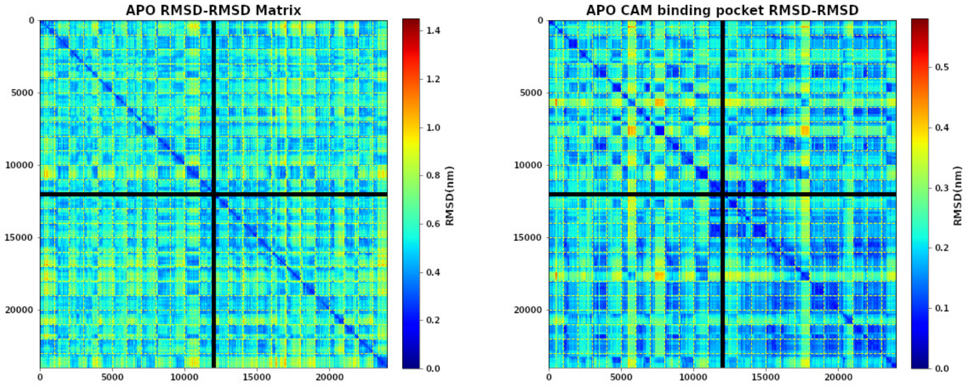


Figure S3: Apo simulations: pairwise RMSDs for the combined 12 replicas using all the protein residue C$\alpha$ atoms (left) or C$\alpha$ atoms of the CAM binding pocket residues (right).

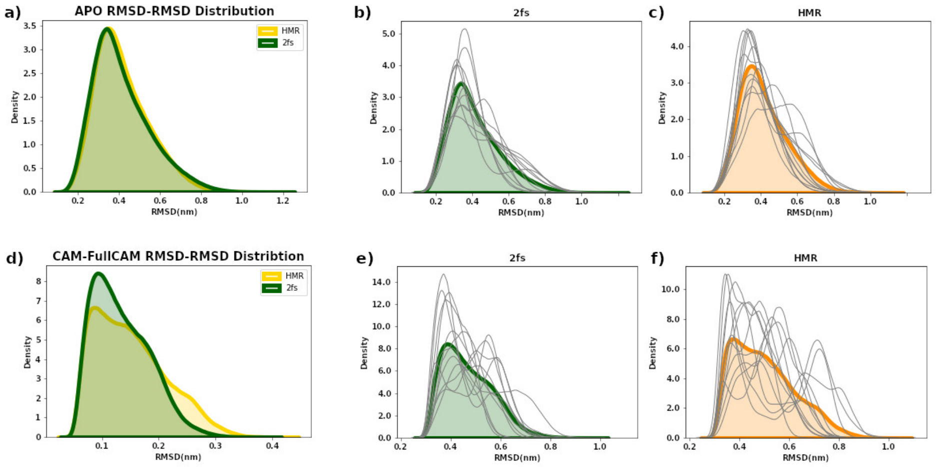


Figure S4: Distribution of all-to-all RMSD values from the apo tetramer simulations comparing data from the HMR to 2fs timestep simulations. Top row (panels a-c), RMSD computed using all protein C$\alpha$ atoms. Lower row (panels d-f), analogous results using the CAM C$\alpha$ atoms of the binding site residues. Panels a) and d) report the combined replica RMSD distributions while the middle and right columns are the 2fs (panels b and e, with combined replicas: green lines) and HMR (panels c and f, with combined replicas: orange lines) respectively. Individual replicas are grey, while the combined replicas are a thick green (2fs) or orange (HMR). Each distribution is individually normalized for visualization.

#### AT-130 HMR/2fs Simulations

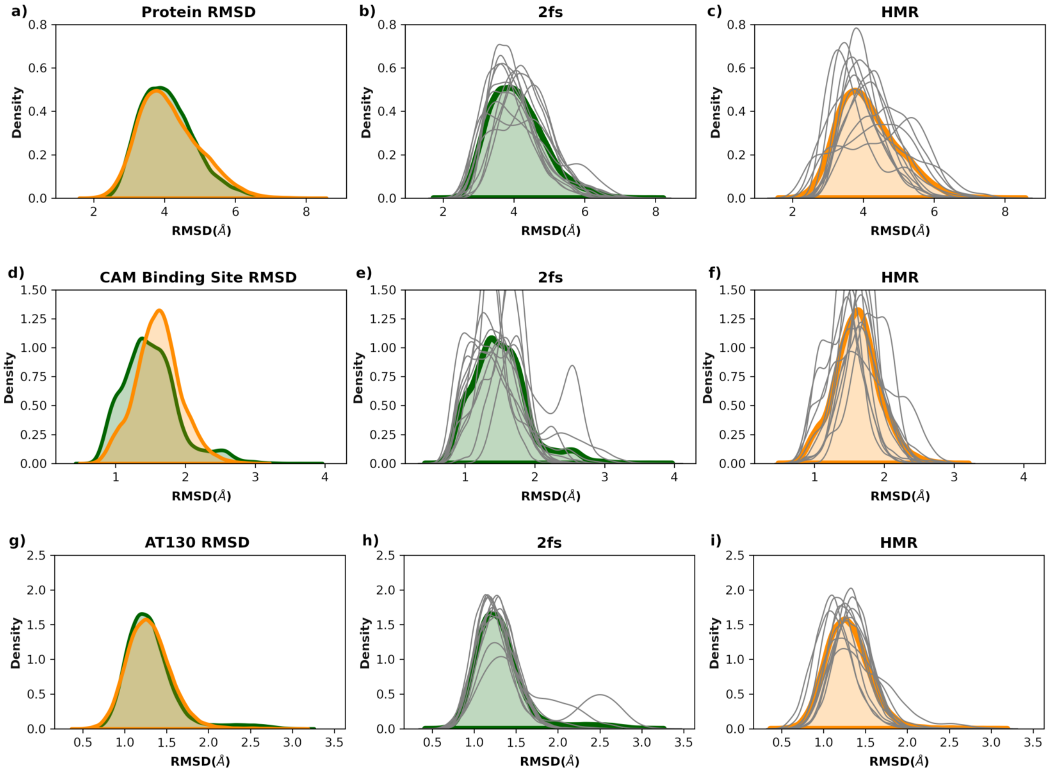


Figure S5: RMSD distributions to the starting structure from the AT-130 bound simulations. Top row (panels a-c): RMSD aligned and calculated using all protein C$\alpha$ atoms. Middle row (panels d-f): RMSD aligned and calculated using the C$\alpha$ atoms of the CAM binding site residues. Lower row (panels g-i): RMSD of the AT-130 heavy atoms after alignment using the C$\alpha$ atoms of the CAM binding site residues. The left column (panels a, d, and g) reports the combined replica RMSD distributions while the middle and right columns are the 2fs (panels b, e, and h with combined replicas: green lines) and HMR (panels d, f, and i with combined replicas: orange lines) respectively, with individual replica data plotted separately (grey lines). Note, for clarity the replicas are separately normalized to 1.

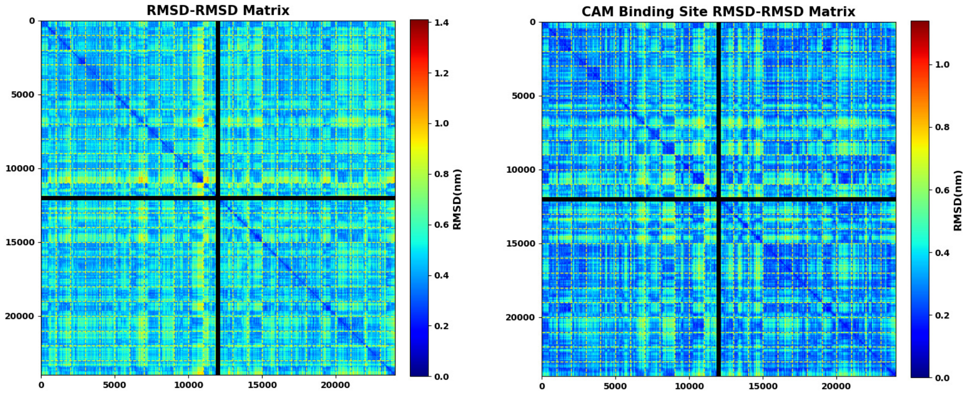


Figure S6: AT-130 bound simulations: pairwise RMSDs using all protein C$\alpha$ atoms (left) or C$\alpha$ atoms of the CAM binding pocket residues (right).

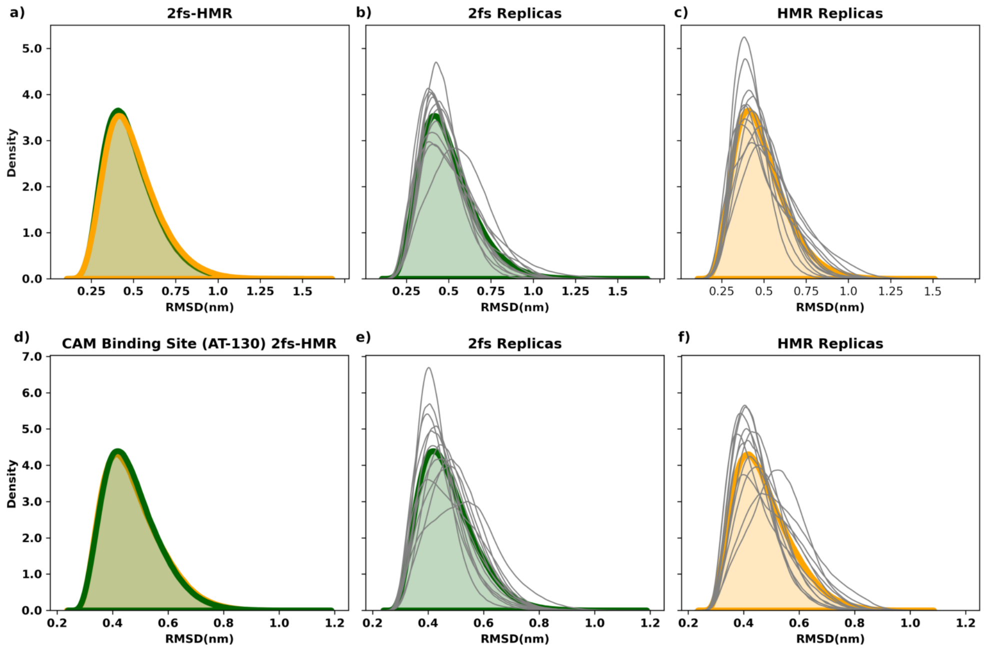


Figure S7: Distribution of all-to-all RMSD values from the AT-130 bound tetramer simulations comparing the HMR to 2fs timestep data. Top row (panels a-c), RMSD computed using all protein C$\alpha$ atoms. Lower row (panels d-f), analogous results using the CAM C$\alpha$ atoms of the binding site residues. Panels a) and d) report the combined replica RMSD distributions while the middle and right columns are the 2fs (panels b and e, with combined replicas: green lines) and HMR (panels c and f, with combined replicas: orange lines) respectively. Individual replicas are grey, while the combined replicas are a thick green (2fs) or orange(HMR). Each distribution is individually normalized for visualization.

### GLS4 HMR/2fs MD Simulations

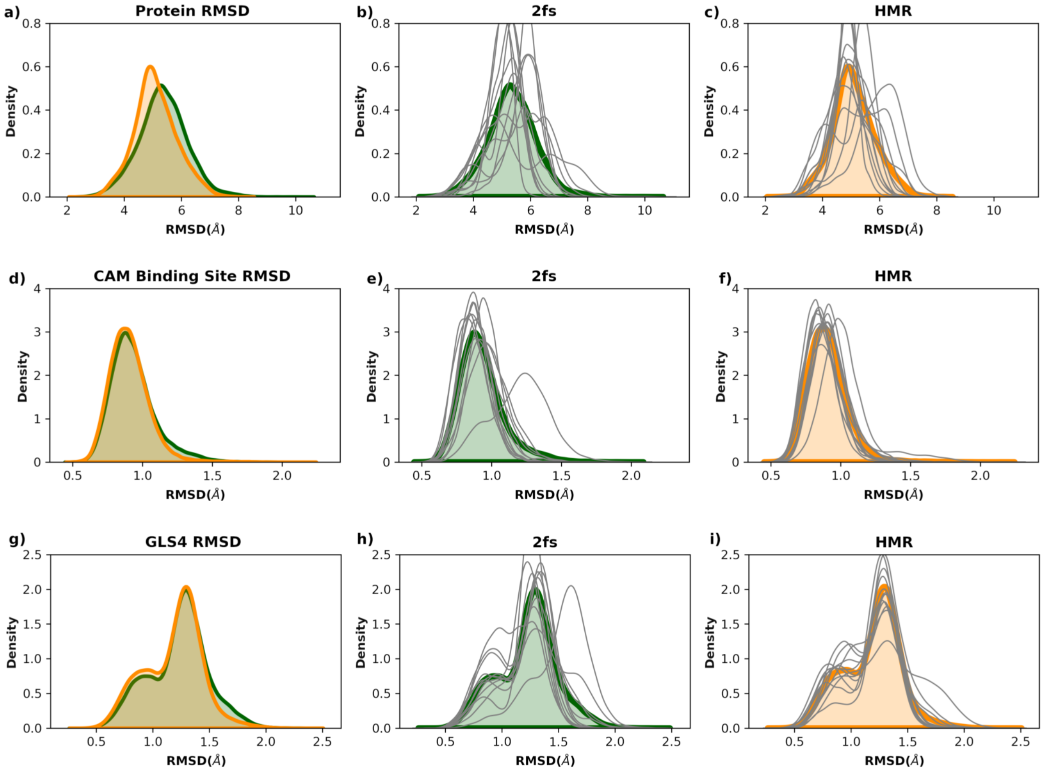


Figure S8: RMSD distributions to the starting structure from the GLS4 bound simulations. Top row (panels a-c): RMSD aligned and calculated using all protein C$\alpha$ atoms. Middle row (panels d-f): RMSD aligned and calculated using the C$\alpha$ atoms of the CAM binding site residues. Lower row (panels g-i): RMSD of the GLS4 heavy atoms after alignment using the C$\alpha$ atoms of the CAM binding site residues. The left column (panels a, d, and g) reports the combined replica RMSD distributions while the middle and right columns are the 2fs (panels b, e, and h with combined replicas: green lines) and HMR (panels d, f, and i with combined replicas: orange lines) respectively, with individual replica data plotted separately (grey lines). Note, for clarity the replicas are separately normalized to 1.

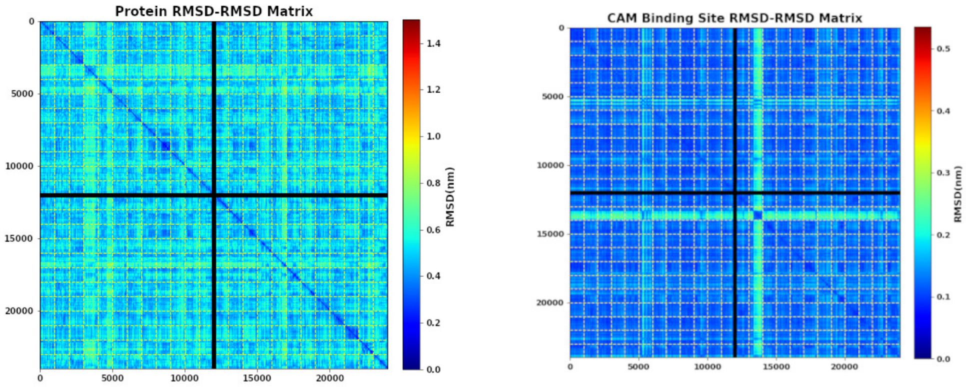


Figure S9: GLS4 bound simulations: pairwise RMSDs using all protein C$\alpha$ atoms (left) or C$\alpha$ atoms of the CAM binding pocket residues (right).

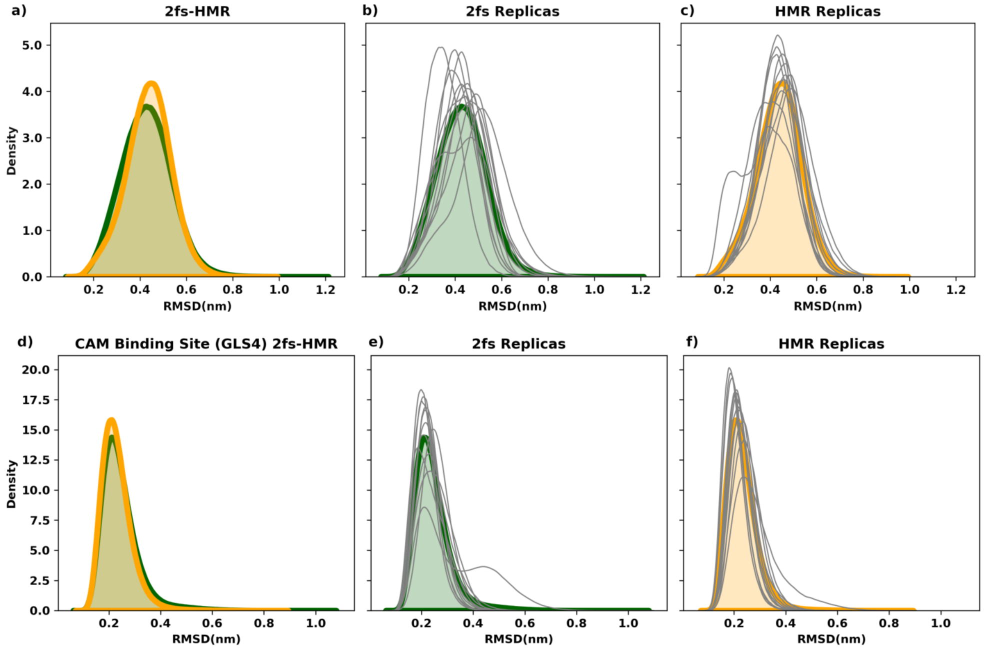


Figure S10: Distribution of all-to-all RMSD values from the GLS4 bound tetramer simulations comparing the HMR to 2fs timestep data. Top row (panels a-c), RMSD computed using all protein C$\alpha$ atoms. Lower row (panels d-f), analogous results using the CAM C$\alpha$ atoms of the binding site residues. Panels a) and d) report the combined replica RMSD distributions while the middle and right columns are the 2fs (panels b and e, with combined replicas: green lines) and HMR (panels c and f, with combined replicas: orange lines) respectively. Individual replicas are grey, while the combined replicas are a thick green (2fs) or orange (HMR). Each distribution is individually normalized for visualization.

#### Comparing 2-fs and HMR results for base and spike angles

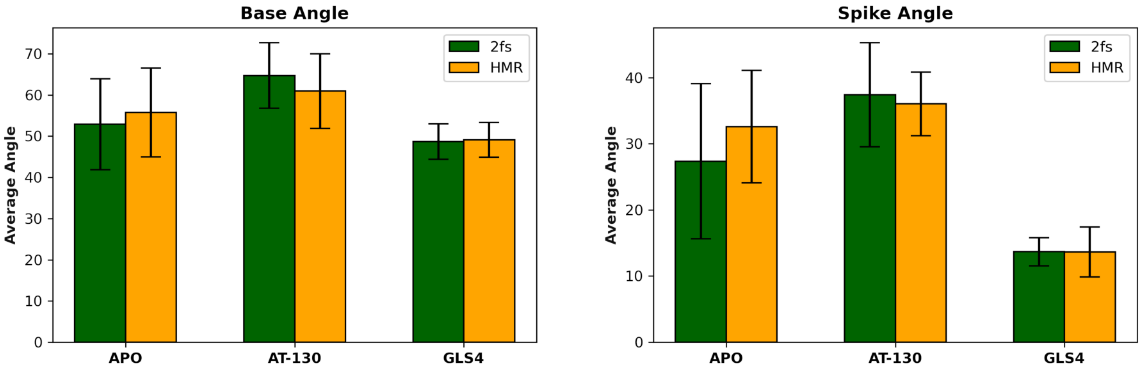


Figure S11: Average base and spike angles with the 95%CI. No statistical difference is seen with the use of hydrogen mass repartitioning.

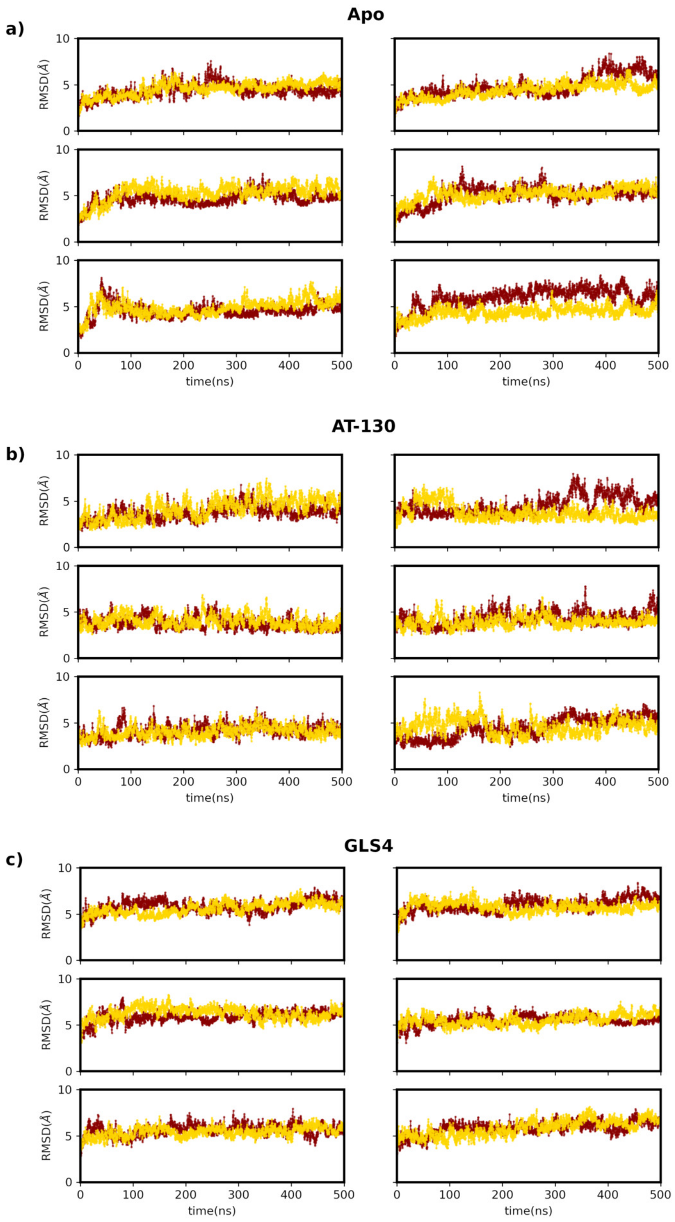


Figure S12: Protein RMSD time-series for the 12 replicas for the a) Apo, b)AT-130, and c) GLS4 simulations. Individual replicas are distinguished by color.

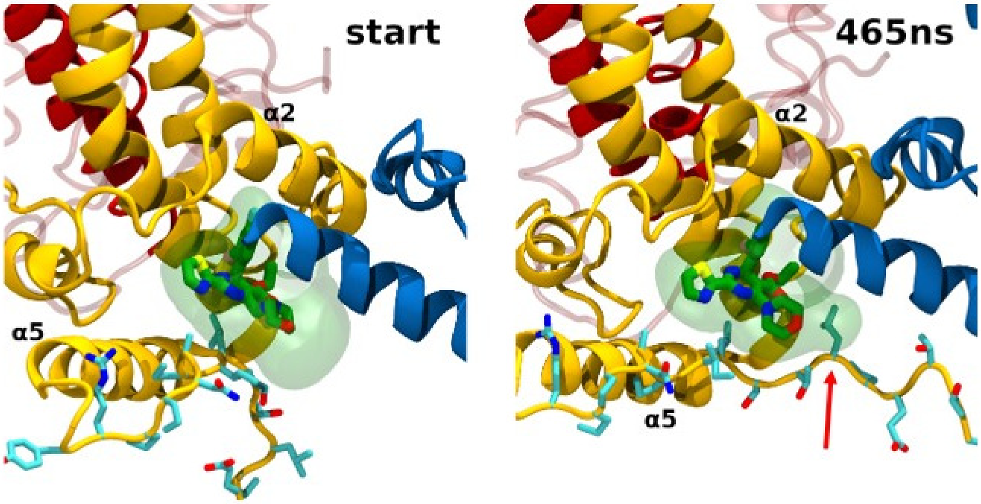


Figure S13: GLS4 Tail Conformations. On left is a snapshot of the starting conformation and right panel is taken at 465ns. The C-terminal tail is rendered licorice and occasionally wraps around and occludes available space near the ligand reducing the volume (area indicated by red arrow).

### WE simulations

#### Apo

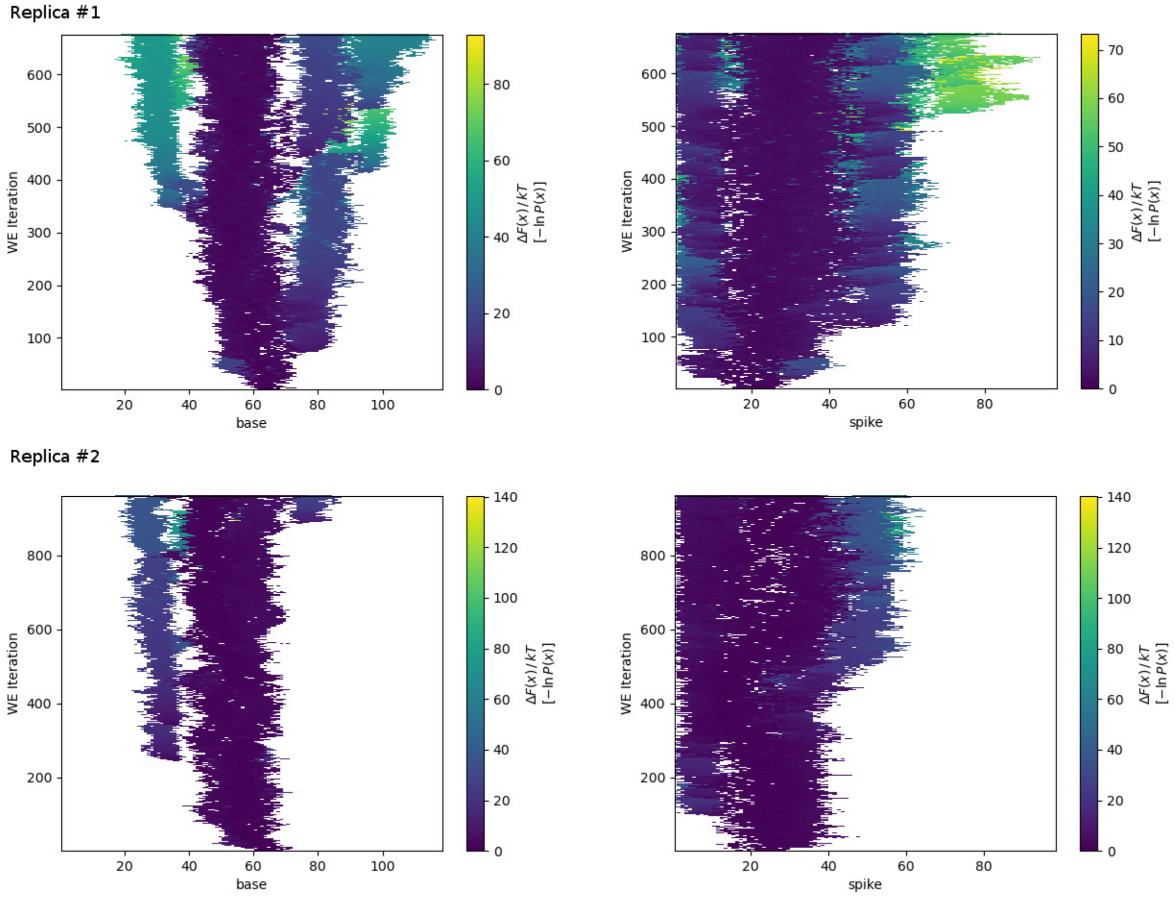


Figure S14: Evolution of the progress coordinates for the two independent Apo replicas at $3\mu$s, with the base angle in the left column and the spike angle in the right column.

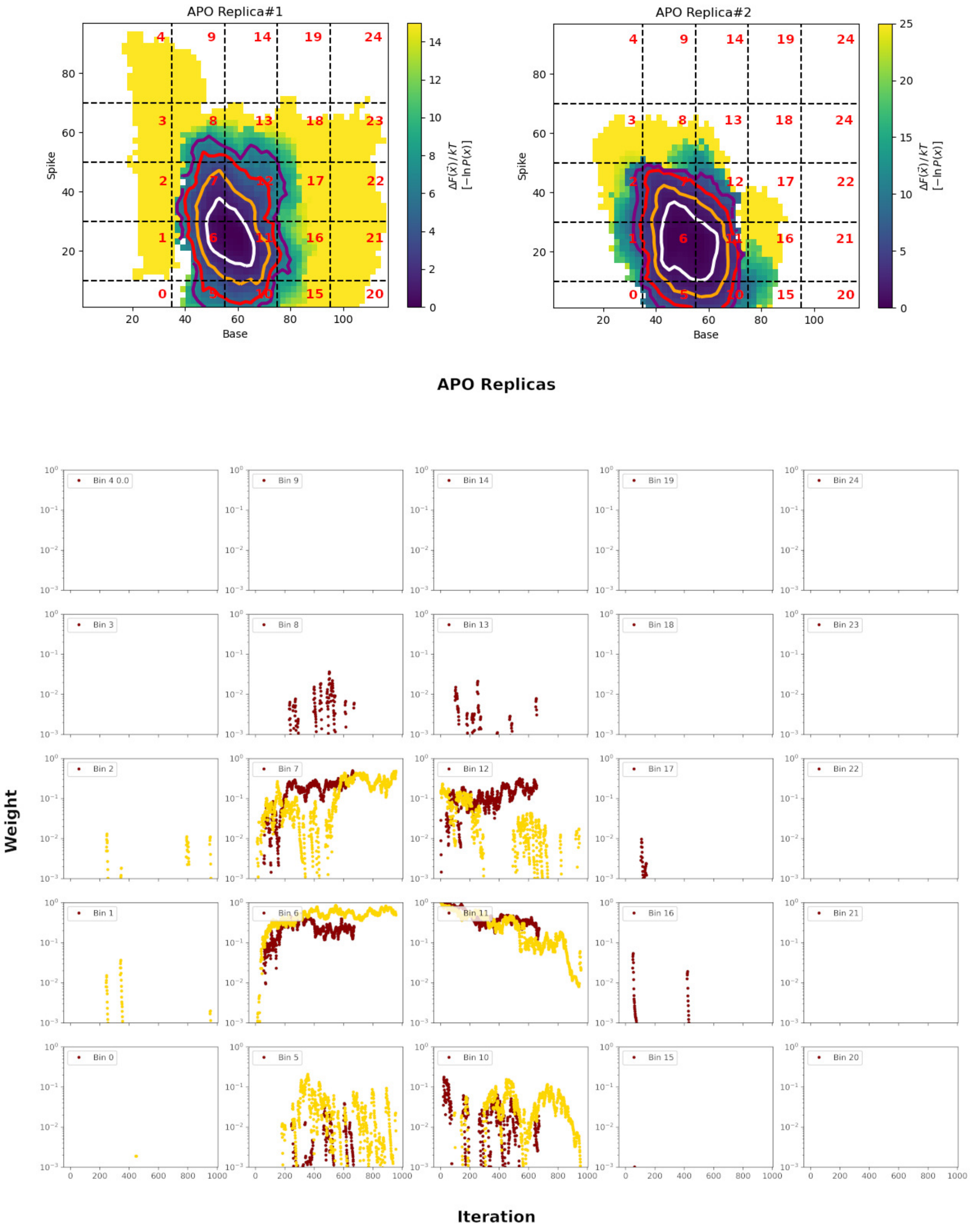


Figure S15: Top row: 2-D progress coordinate landscape with bins labeled for the two replicas. Contours are plotted at 1kT(white), 2.5kT(orange), 5.0kT(red), 7.5kT(purple). Lower panel: Bin weights as a function of WE iteration, with the two replicas colored red (replica #1) or gold (replica#2).

## AT-130

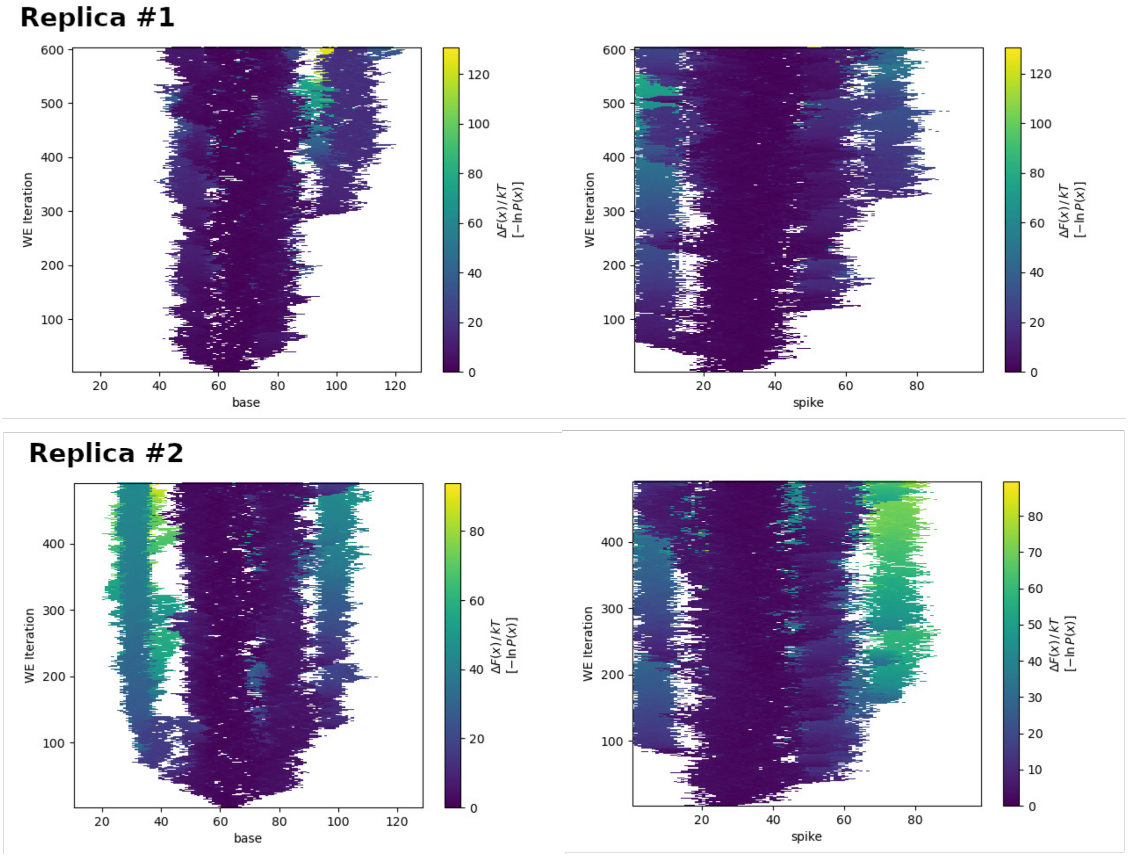


Figure S16: Evolution of the progress coordinates for the two independent AT-130 replicas at 3$\mu$s, with the base angle in the left column and the spike angle in the right column.

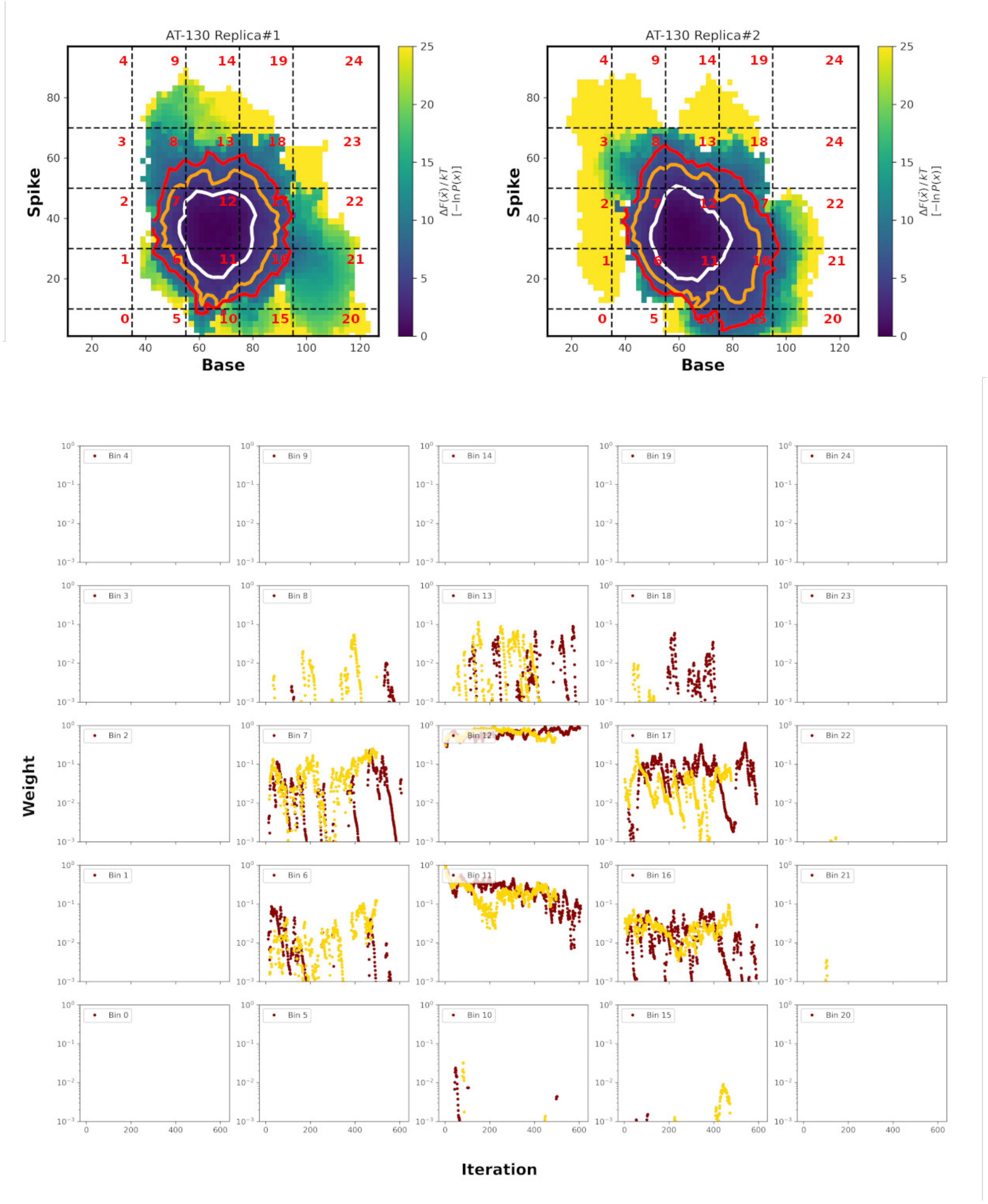


Figure S17: AT-130 Simulations: top row: 2D progress coordinate landscape with bins labeled for the two replicas. Contours are plotted at 2.5kT(white), 5kT(orange), 7.5kT(red). Lower panel: Bin weights as a function of WE iteration, with the two replicas colored red (replica #1) or gold (replica#2).

#### GLS4

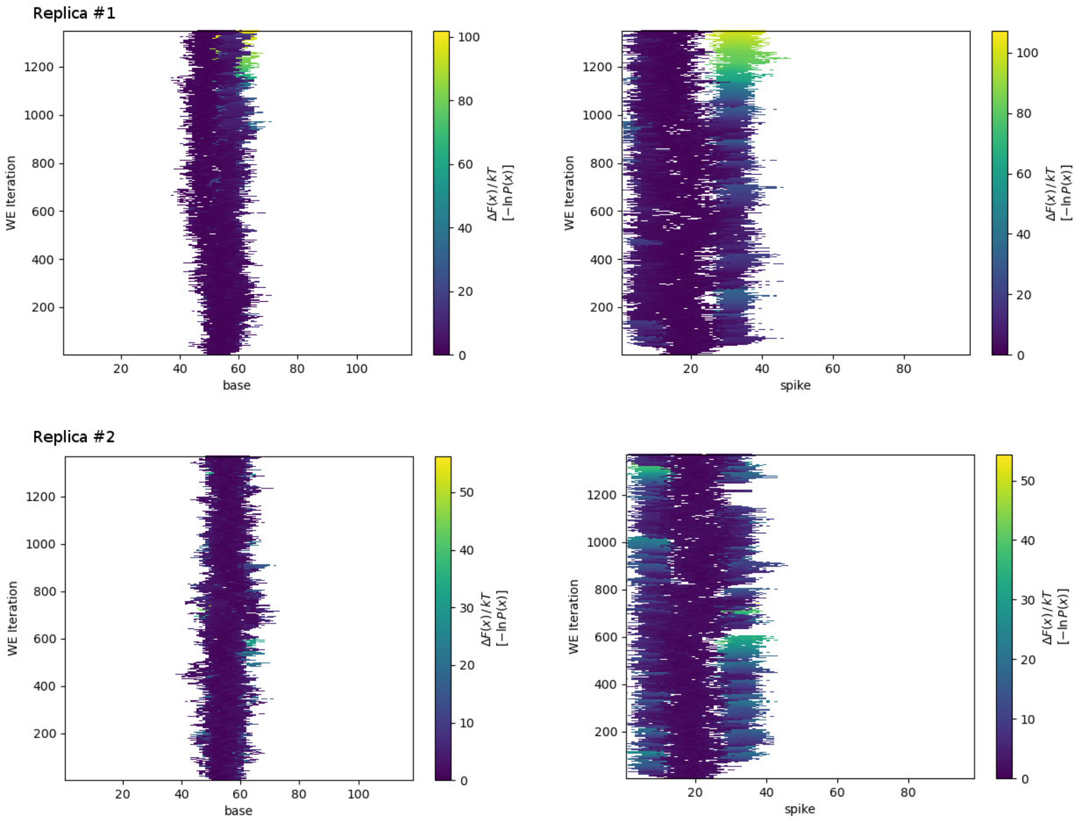


Figure S18: Evolution of the progress coordinates for the two independent GLS4 replicas, with the base angle in the left column and the spike angle in the right column.

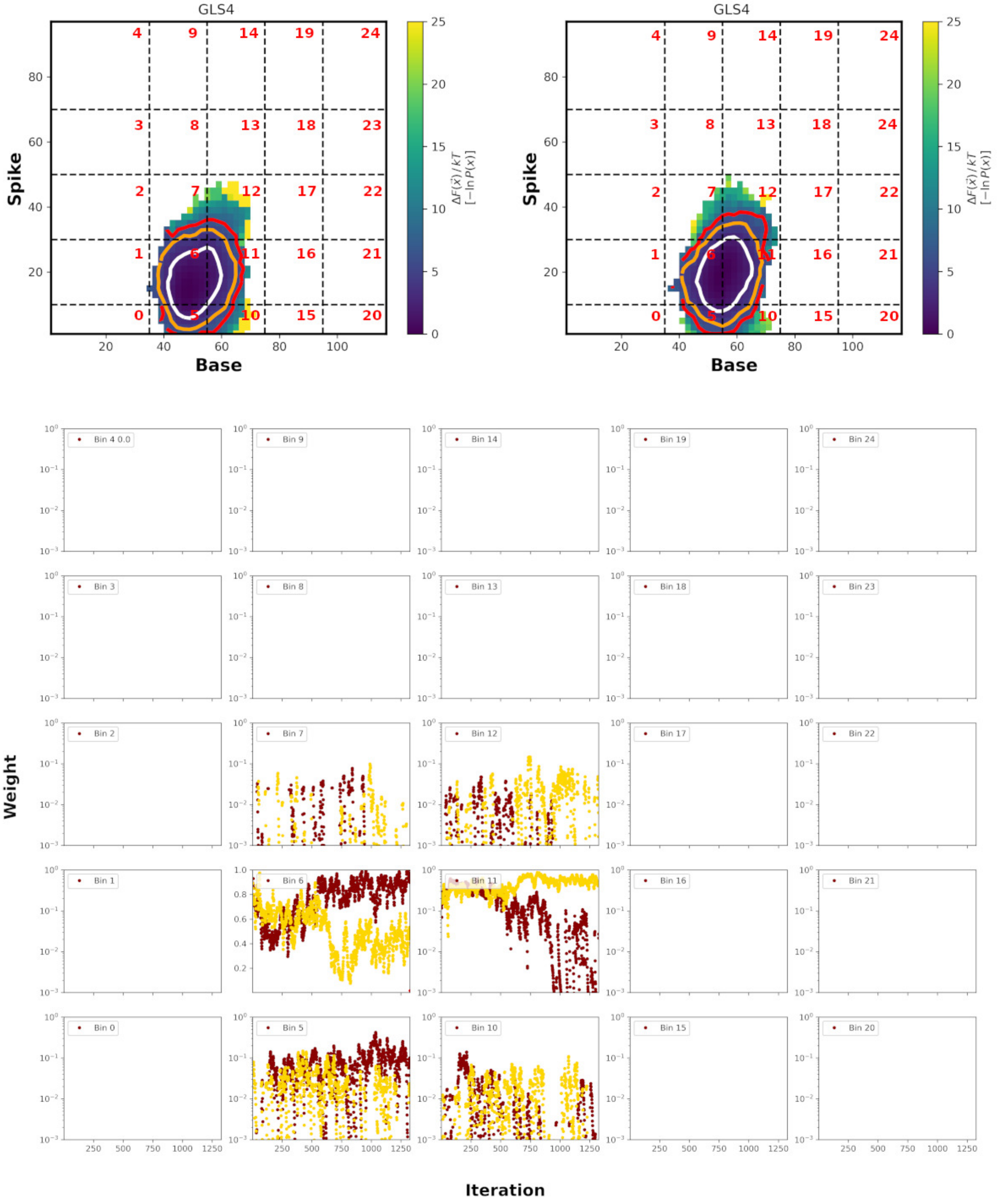


Figure S19: GLS4 Simulations: top row is 2D progress coordinate energy landscape for the two replicas. Contours are plotted at 2.5kT (white), 5.0kT(orange), 7.5kT(red). Lower panel: Bin weights as a function of WE iteration, with the tow replicas colored red (replica 1) or gold (replica 2).

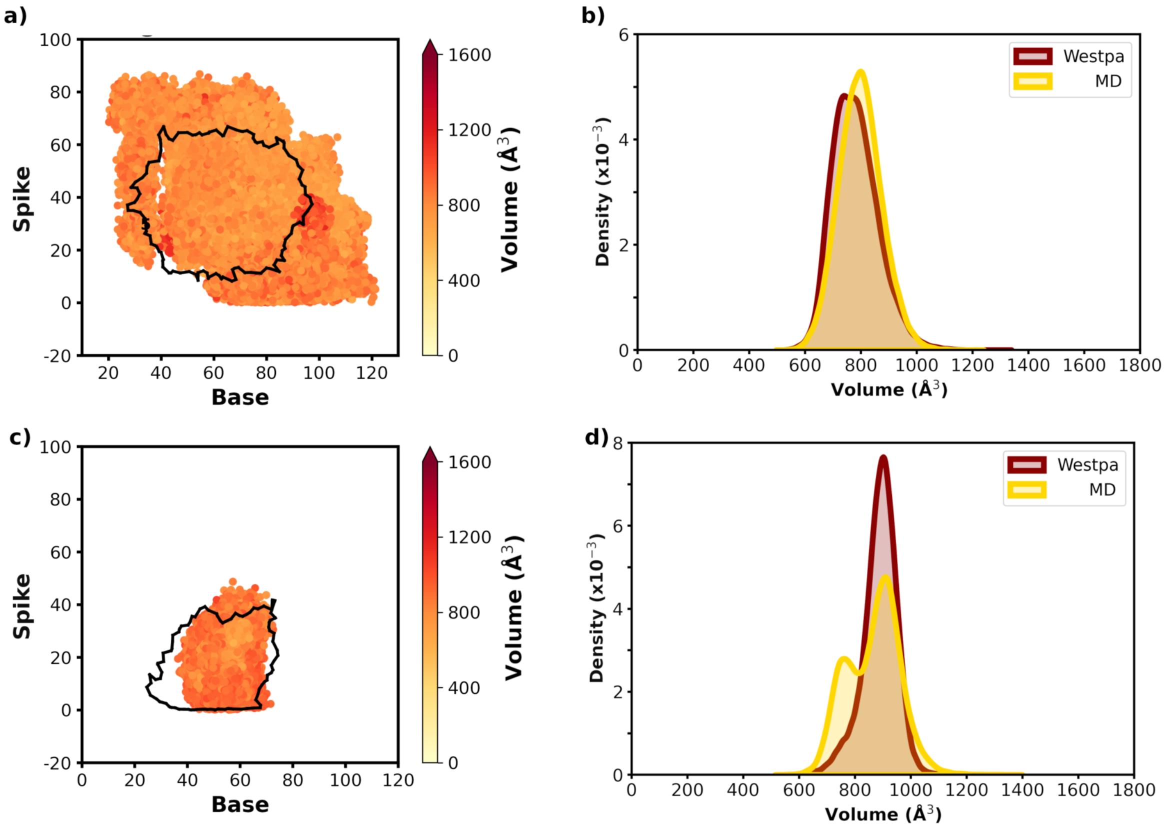


Figure S20: Base and spike angle distributions as well as CAM binding pocket volumes for the AT-130 and GLS4 WE and standard MD. Scatter plots of the base and spike angle progress coordinates (unweighted) for the AT-130 (panel a) and the GLS4 (panel c) simulations. Collective data is reported, with the first 250 iterations of WE simulation removed and the first 10ns of cMD data removed. For clarity, the cMD data is rendered as a boundary (black line) which outlines the region sampled by these 12 replicas, while the WE data is a scatter plot colored by the CAM pocket volume, calculated using Epock. Panels b) and d), histograms of the WE and standard MD derived ligand binding pocket volumes for the AT-130 (panel b) and GLS4 (panel d) simulations.

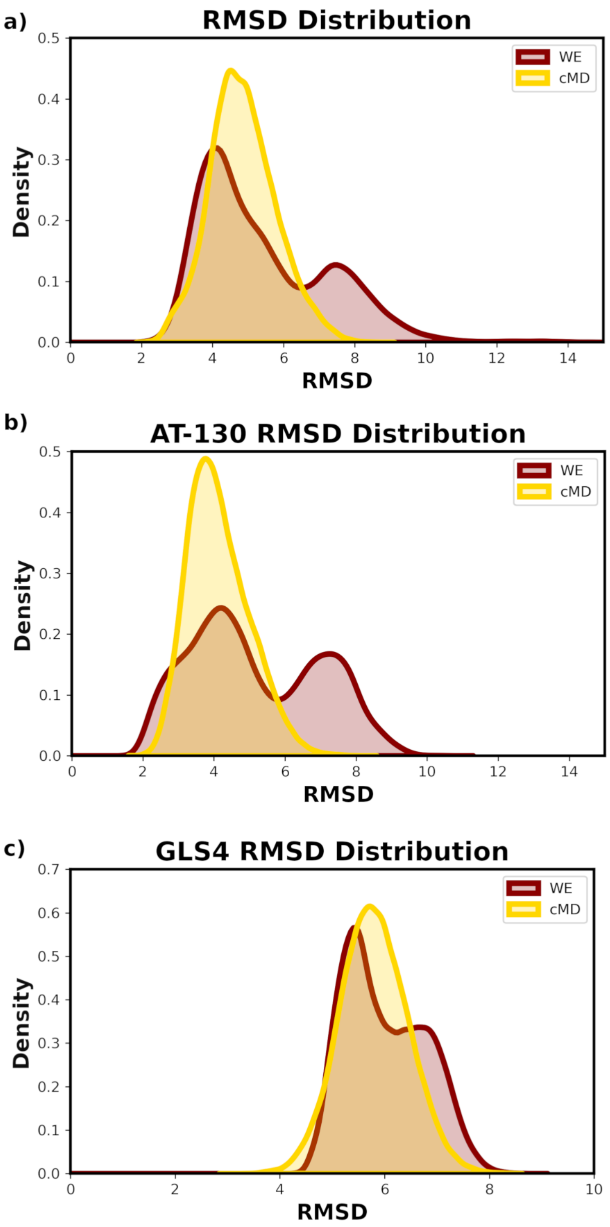


Figure S21: Protein RMSD Distributions for WE(darkred, unweighted data) and standard (gold) a) Apo simulations, b) AT-130 simulations and c) GLS4 simulations.

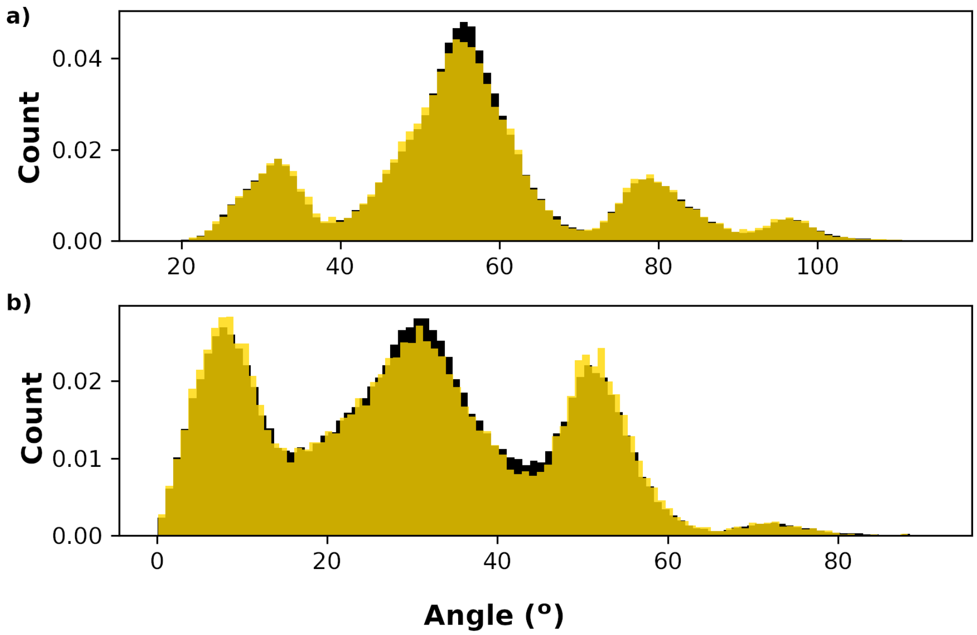


Figure S22: Base (top panel a) and spike (lower panel b) histograms of WE (unweighted) downsampled data. The black histograms are the full data set, while the gold are downsampled, with every 4th data point retained).

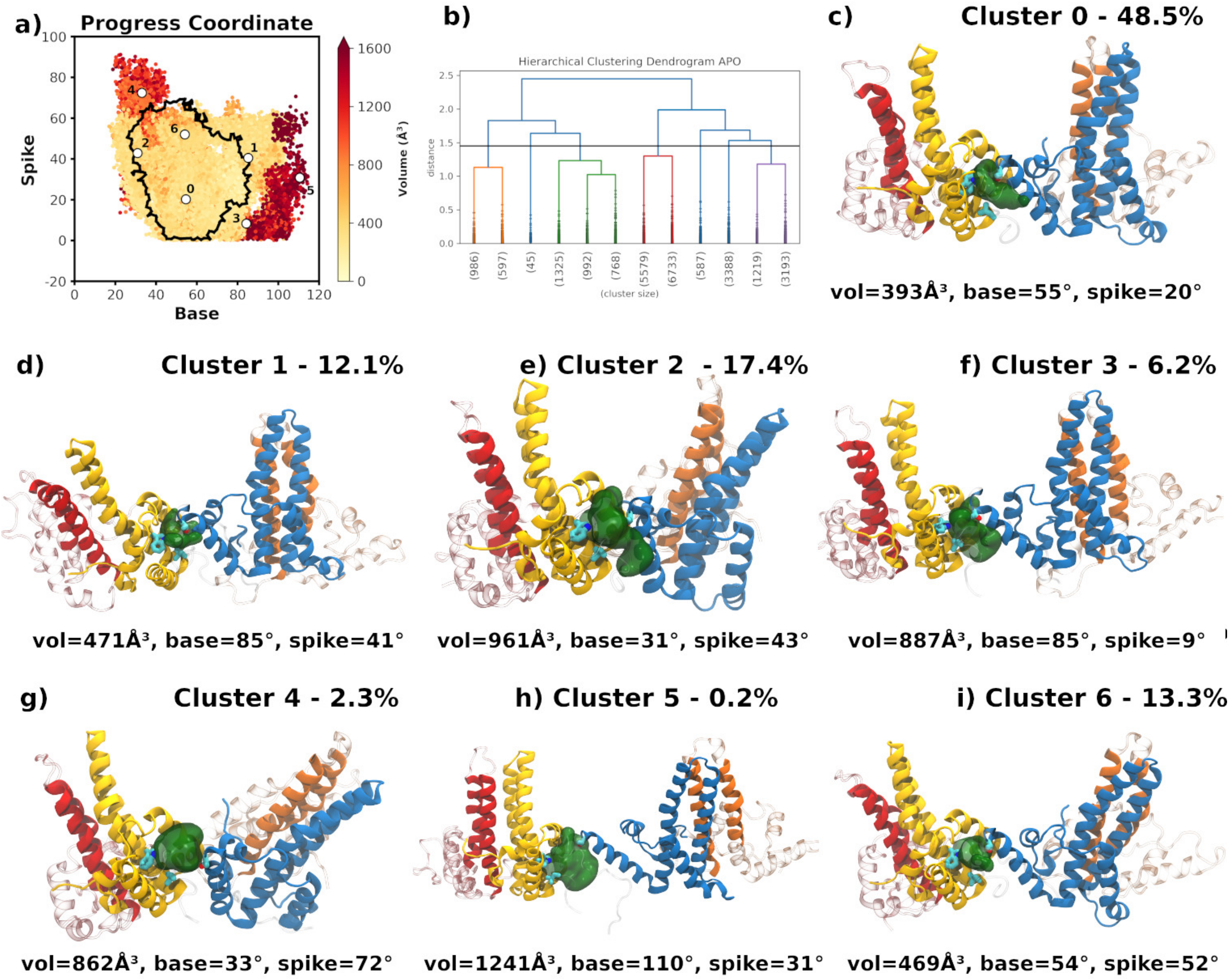


Figure S23: Clustering of the Apo WE simulations. a) Scatter plot of volumes for the base and spike angle progress coordinates (unweighted; see Fig. 5a in main text). b) Hierarchical Dendrogram for the average euclidean clustering. Solid line represents the cut-off for 7 clusters. Panels c) - i) representative conformations, taken as the structure with base/spike angle pair closest to the center, from each cluster. The structures are rendered as chain A (red), chain B (gold), chain C (blue), and chain D (orange), with the Epock volumes illustrated with a dark green surface. The volume, base and spike angles are reported below each cluster and the percentage of total conformations contributed by each cluster is indicated.

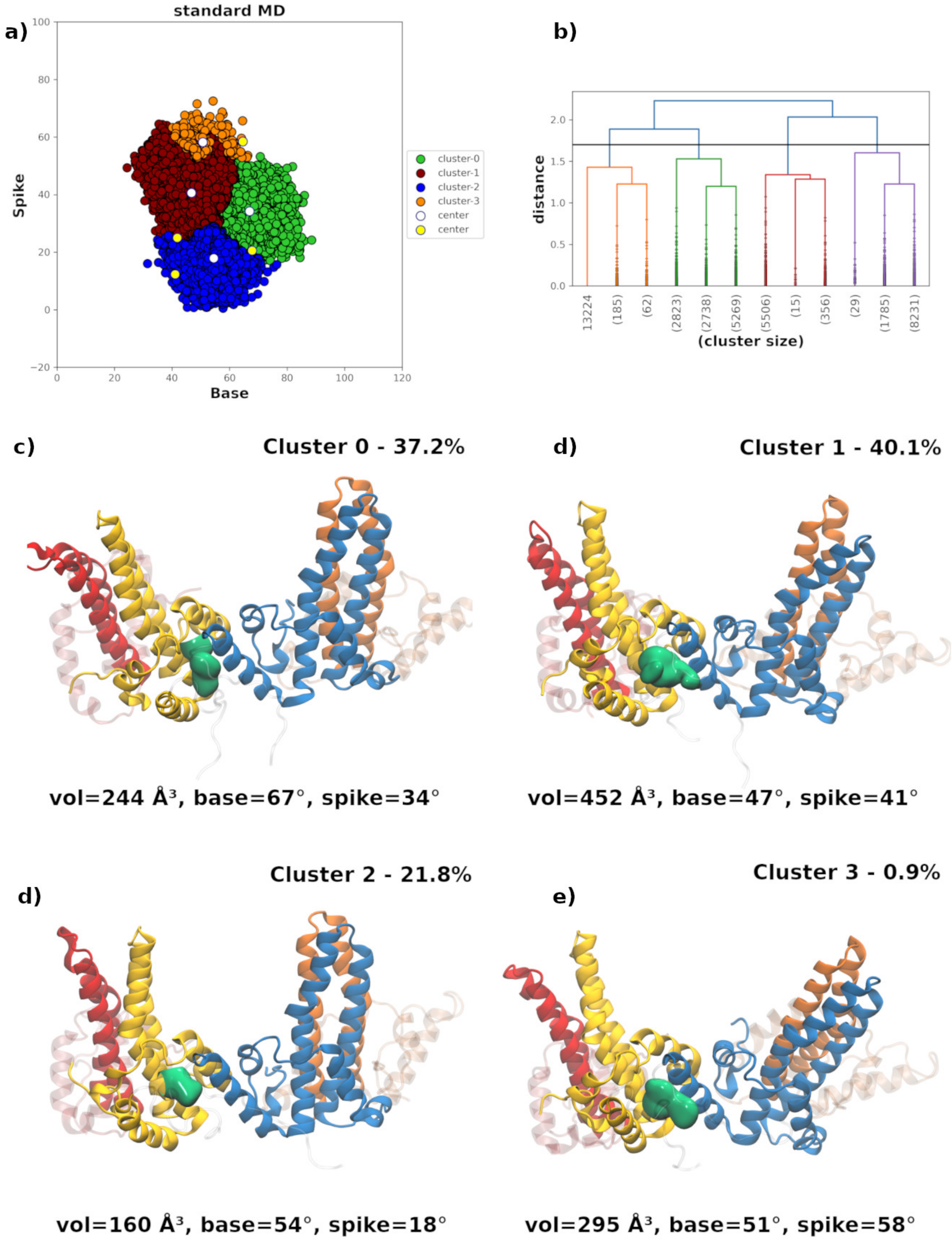


Figure S24: Clustering of the Apo standard MD simulations. a) Scatter plot of the base/spike angles colored by cluster membership. The conformation in each cluster closest to the cluster center is indicated with a white circle. Yellow circles represent alternate choices (see Figure S19). b) Hierarchical Dendrogram for the average euclidean clustering of the Apo MD, with initial 10ns removed. Solid line represents the cut-off for 4 clusters. Panels c) - f) representative conformations, taken as the structure with base/spike angle pair closest to the center, from each cluster. The structures are rendered as chain A (red), chain B (gold), chain C (blue), and chain D (orange), with the Epock volumes illustrated with a dark green surface. The volume, base and spike angles are reported below each cluster and the percentage of total conformations contributed by each cluster is indicated.

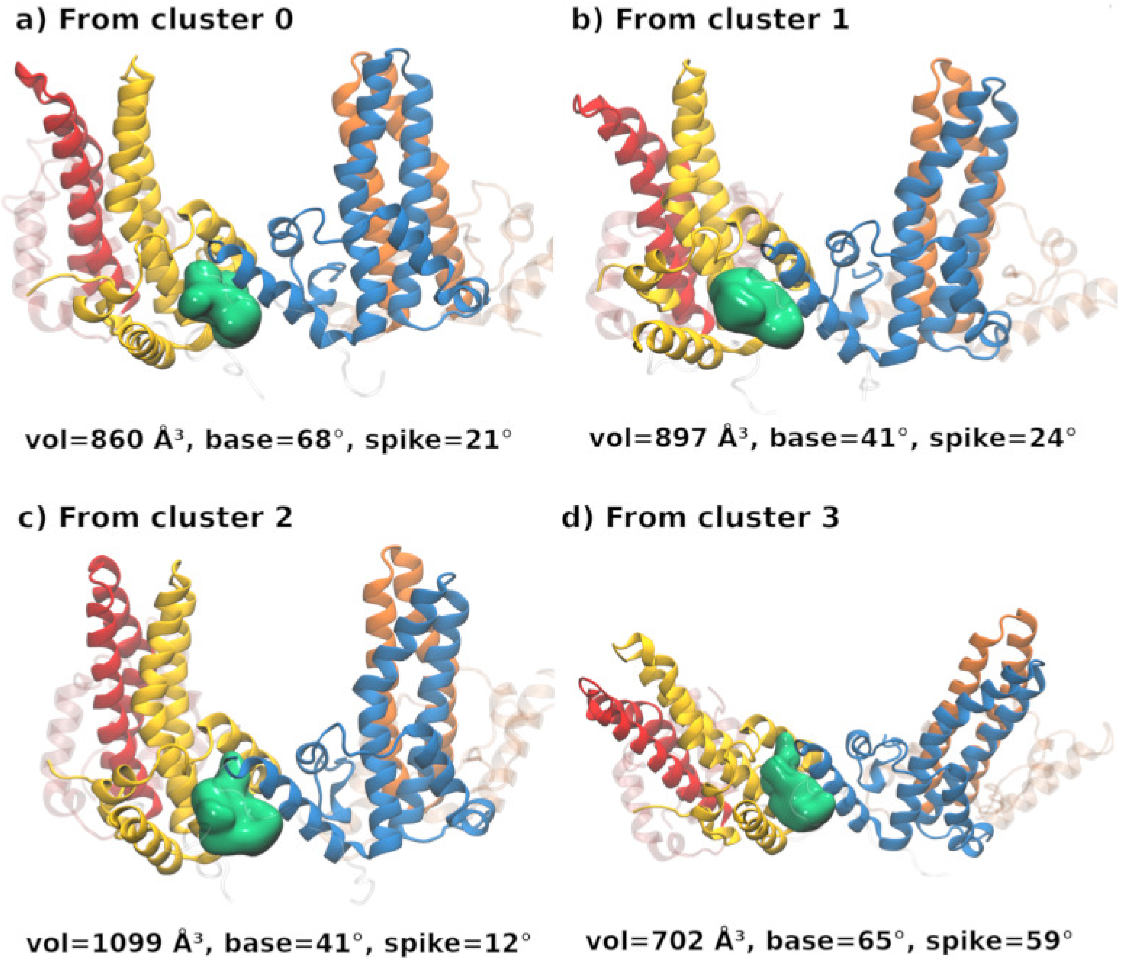


Figure S25: Clustering of the Apo standard MD simulations: structures with large volumes. Panels a) - d) Conformations were taken from each cluster as follows: The 5% percent with the largest distance from the cluster center were sorted by binding pocket volume, and the conformation with the largest volume was extracted. The base/spike values are rendered in Figure SI24a as the yellow circles. The structures are rendered as chain A (red), chain B (gold), chain C (blue), and chain D (orange), with the Epock volumes illustrated with a dark green surface. The volume, base and spike angles are reported below each cluster and the percentage of total conformations contributed by each cluster is indicated.

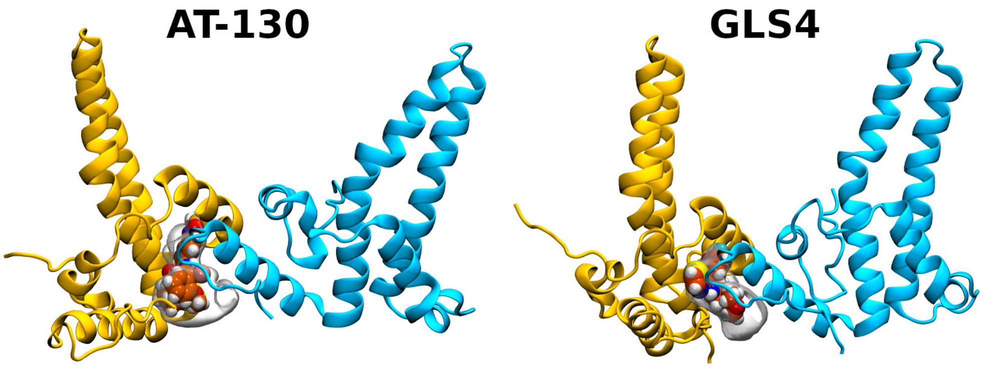


Figure S26: Illustration of the a) AT-130 and b) GLS4 CAM binding pocket and the rendered Epock volume. Chain B (gold) and Chain C (blue) are rendered in ribbons, as in Figure 1. Chains A and D are not displayed. The ligand is rendered in VDW spheres and the Epock volume is rendered as a transparent white surface indicating that the region used in the Epock volume calculation is reasonable.
